# Transposable element-mediated co-option drives the evolution of the miRNA regulatory system in *Oryza* AA-genome species

**DOI:** 10.64898/2026.01.18.700148

**Authors:** Yun Zhang, Richard S. Taylor, Li-Ping Zhang, Philip Donoghue, Li-Zhi Gao

## Abstract

MicroRNAs (miRNAs) are key post-transcriptional regulators in plants, yet the evolutionary dynamics of the entire miRNA-mediated regulatory system remain poorly understood. We performed small RNA sequencing and comparative genomics analyses across all eight AA-genome species of *Oryza* within a well-resolved phylogenetic framework. We found that miRNA gene (MIRNA) families evolve with low birth-death rates, while individual paralogous MIRNAs turn over rapidly. We identified a novel mechanism in which transposable element (TE) insertions *de novo* generates functional miRNA target sites. miRNA target genes are characterized by distinct evolutionary signatures, including greater sequence length, lower GC content, moderately highly expression, reduced expression variability, and higher evolutionary conservation, which is consistent with their role as a conserved kernel regulatory subsystem. In contrast, the majority of recently evolved target genes that have acquired their miRNA binding sites through TE insertions tend to exhibit the opposite set of features. Furthermore, we uncovered co-evolutionary signatures between duplicated MIRNAs and their target genes, as well as between MIRNAs and phased siRNAs (phasiRNAs). Taken together, our study reveals multi-level evolutionary dynamics driven by TEs that rapidly generate new regulatory circuit and proposes a generalizable TE-MIRNA co-option model for regulatory network expansion in plants.

## INTRODUCTION

MicroRNAs (miRNAs) are a class of small, endogenous noncoding RNAs that are produced by Dicer-catalyzed excision from hairpin structure precursors transcribed from MIRNA genes (MIRNAs). The majority of miRNAs are able to negatively regulate gene expression via direct RNA-induced silencing complex (RISC) binding to target mRNAs, resulting in the transcript degradation or translational repression, while small fractions have developed specific properties that regulate other transcriptional or posttranscriptional silencing pathways (Yu, et al. 2026). miRNAs are pivotal regulators of gene expression in plants, influencing diverse processes from development to environmental adaptation, and therefore, are regarded as important candidates for bioengineering to improve crop yields and food security (Ding, et al. 2020, Sang, et al. 2023, Fahad, et al. 2025).

While the functional roles of individual miRNAs are increasingly well-characterized, the evolutionary forces and mechanisms that shape the entire miRNA-mediated regulatory system — comprising the MIRNAs themselves, their target sites, and the resulting network interactions — remain a fundamental and unresolved question in evolutionary genomics. Understanding how MIRNAs, their targets, and their interactions evolve is crucial not only for deciphering regulatory network evolution but also for harnessing miRNAs in crop improvement (D’Ario et al., 2017). Prevailing views on the rapid birth-death rate of plant MIRNAs may stem from limited phylogenetic sampling and inconsistent annotation practices (Fahlgren et al., 2007; Meng et al., 2012; Taylor et al., 2014). While the inverted duplication model elucidates certain *de novo* origins of MIRNAs (Allen et al., 2004), and subsequent work proposed that transposable element (TE) insertions could generate novel miRNA target sites (Zhang et al., 2011), the mechanisms driving target-site emergence—a key process in regulatory network expansion—are still not fully resolved. Although miRNA target genes have been reported to exhibit lower polymorphism levels than genomic averages (Takuno and Innan, 2011), a systematic analysis of their broader evolutionary features—such as gene structure, GC content, expression patterns, selective pressure, duplication history, and evolutionary age—is still lacking. Furthermore, while miRNAs may co-evolve with their targets, direct evidence in plants remains sparse (Tang et al., 2010; Zhang et al., 2011; Wang, et al. 2024). miRNAs can also trigger phased siRNAs (phasiRNAs) from *PHAS* loci, producing secondary silencing signals that influence regulatory networks (Fei et al., 2013; Lian, et al. 2025); however, how such secondary pathways co-evolve with miRNAs is largely unknown.

A major obstacle to deciphering the evolution of this regulatory system has been the absence of both accurate MIRNA annotation and comprehensive evolutionary analysis of target genes within a well-resolved phylogenetic framework. To date, complete MIRNA repertoires have been described in only one pair of close relatives (*Arabidopsis thaliana* and *A. lyrata*) (Fahlgren et al., 2010; Ma et al., 2010), and comparative studies of miRNA target-gene evolution remain preliminary (Takuno and Innan, 2011). In rice (*Oryza sativa*), changes in the miRNA regulatory system have been linked to phenotypic variation (Liu et al., 2013; Wang et al., 2012; Zhang et al., 2014; Rong, et al. 2024), highlighting its evolutionary significance. To address these gaps, we conducted a genome-wide comparative analysis of miRNAomes and transcriptomes across all eight AA-genome species in the genus *Oryza*—including two cultivated and six wild rice species—within a definitive phylogenetic context. This study provides systematic insights into the evolutionary mechanisms shaping the miRNA-mediated regulatory system in plants.

## RESULTS

### Divergent evolutionary dynamics between conserved MIRNA families and rapidly evolving individual MIRNAs

We performed deep sequencing of small RNAs from 30-day-old roots, 30-day-old shoots, panicles at the booting stage, and flag leaves at the booting stage of seven AA-genome *Oryza* species: *O. rufipogon* (RUF), *O. nivara* (NIV), *O. glaberrima* (GLA), *O. barthii* (BAR), *O. glumaepatula* (GLU), *O. longistaminata* (LON), and *O. meridionalis* (MER). Integrated with public small RNA data from *O. sativa* ssp. *japonica* cv. Nipponbare (SAT) using our plant MIRNA annotation pipeline (see Methods), we identified a total of 2,497 individual MIRNAs, ranging from 273 in NIV to 349 in RUF. Based on sequence similarity of mature and precursor sequences, these loci were clustered into 110 MIRNA families. Among these, 42 families are newly reported here, while 68 were previously documented in miRBase v21 (Kozomara and Griffiths-Jones, 2014) (Supplemental Table 1). Homology searches further identified 270 MIRNAs belonging to 49 families in *O. brachyantha* and 120 MIRNAs from 28 families in *Sorghum bicolor*.

To reconstruct the evolutionary history of MIRNAs within the AA-genome *Oryza* species, we inferred the latest evolutionary point of each MIRNA family based on its phylogenetic distribution and mapped these onto a reconstructed *Oryza* phylogeny (Figure 1). Given an estimated divergence time of ∼4.8 million years (Myr) for AA-genome *Oryza* species (Zhang et al., 2014), the acquisition rate was estimated at ∼3.15 MIRNA families per Myr within the AA clade. The acquisition rate of individual MIRNAs was considerably higher, at ∼17.37 MIRNAs per Myr. Subsequent loss rates were estimated at ∼3.23 families and ∼14.53 MIRNAs per Myr, respectively. To investigate factors influencing loss, we categorized MIRNA families into three age groups: ancient (originating before angiosperms), medium (originating after angiosperms but before the genus *Oryza*), and young (originating within *Oryza*). Older MIRNA families showed significantly higher conservation than younger families (Supplemental Figure 1A and 1B), a pattern also observed at the level of individual MIRNAs (Supplemental Figure 1C and 1D).

**Figure 1.**
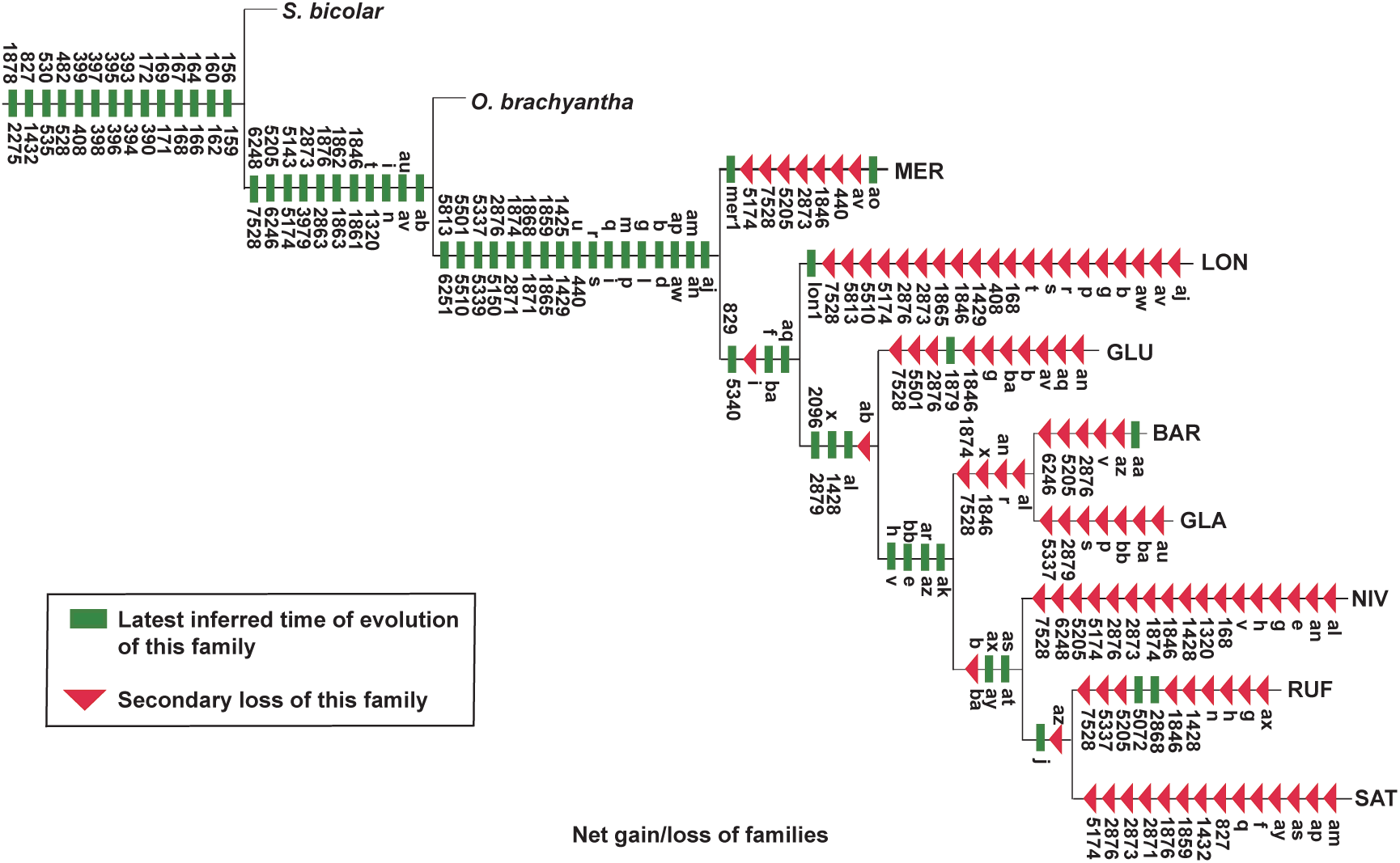
Evolutionary dynamics of MIRNA families in AA-genome *Oryza* species. The most recent evolutionary origin of each MIRNA family was inferred from its phylogenetic distribution across the established *Oryza* phylogeny. *O. brachyantha* and *S. bicolor* were used as outgroups. The divergence time of AA-genome *Oryza* species is estimated at ∼4.8 million years (Zhu et al., 2014).

### Transposable element insertions drive *de novo* origin of miRNA target sites

To understand the evolution of the MIRNA-mediated gene regulatory system—wherein MIRNAs function primarily by interacting with target genes—we investigated the generation of miRNA target sites and the evolutionary dynamics of miRNA target genes, using the AA-genome *Oryza* species phylogeny as our framework. We first examined the relationships among MIRNAs, miRNA target sites, and transposable elements (TEs) across the eight AA-genome *Oryza* species. As summarized in Supplemental Tables 2–3, 19.42% of MIRNAs were associated with TEs, 29.56% of MIRNAs had target sites overlapping TEs, and 20.37% of miRNA target genes contained TE-overlapping miRNA target sites. We further identified MIRNAs whose own sequences and at least one of their target sites overlapped with the same TE. Across the eight species, 16 (SAT), 39 (RUF), 14 (NIV), 15 (GLA), 18 (BAR), 15 (GLU), 30 (LON), and 26 (MER) such MIRNAs were found (Supplemental Table 4).

We manually inspected these MIRNAs and their target sites using a synteny map constructed for the eight AA-genome *Oryza* species with MERCATOR and MAVID (Bray and Pachter, 2004). Notably, osa-miR1868—a reported post-transcriptional regulator involved in rice grain development (Wu et al., 2009; Zhu et al., 2008)—was conserved across all eight species, with its sequence consistently overlapping a *Tourist*-like miniature inverted-repeat transposable element (MITE) (Supplemental Figure 2). In SAT, the target site of osa-miR1868 resides in the fourth exon of LOC_Os01g40320 and overlaps the same MITE (Figure 2). This gene lies within a conserved syntenic region across the eight species (Figure 2; Supplemental Table 5) and is annotated with Gene Ontology terms related to reproduction, post-embryonic development, and embryo development. Frameshift mutations have rendered its orthologs pseudogenes in the NIV, GLA, BAR, and MER genomes (Supplemental Figure 3). The same TE is inserted in the syntenic region of RUF, NIV, GLA, BAR, and GLU, suggesting insertion in their common ancestor. In SAT and GLU, exonization of the inserted TE furnished sequence that later formed the miRNA-binding site on the host gene (Figure 2).

**Figure 2.**
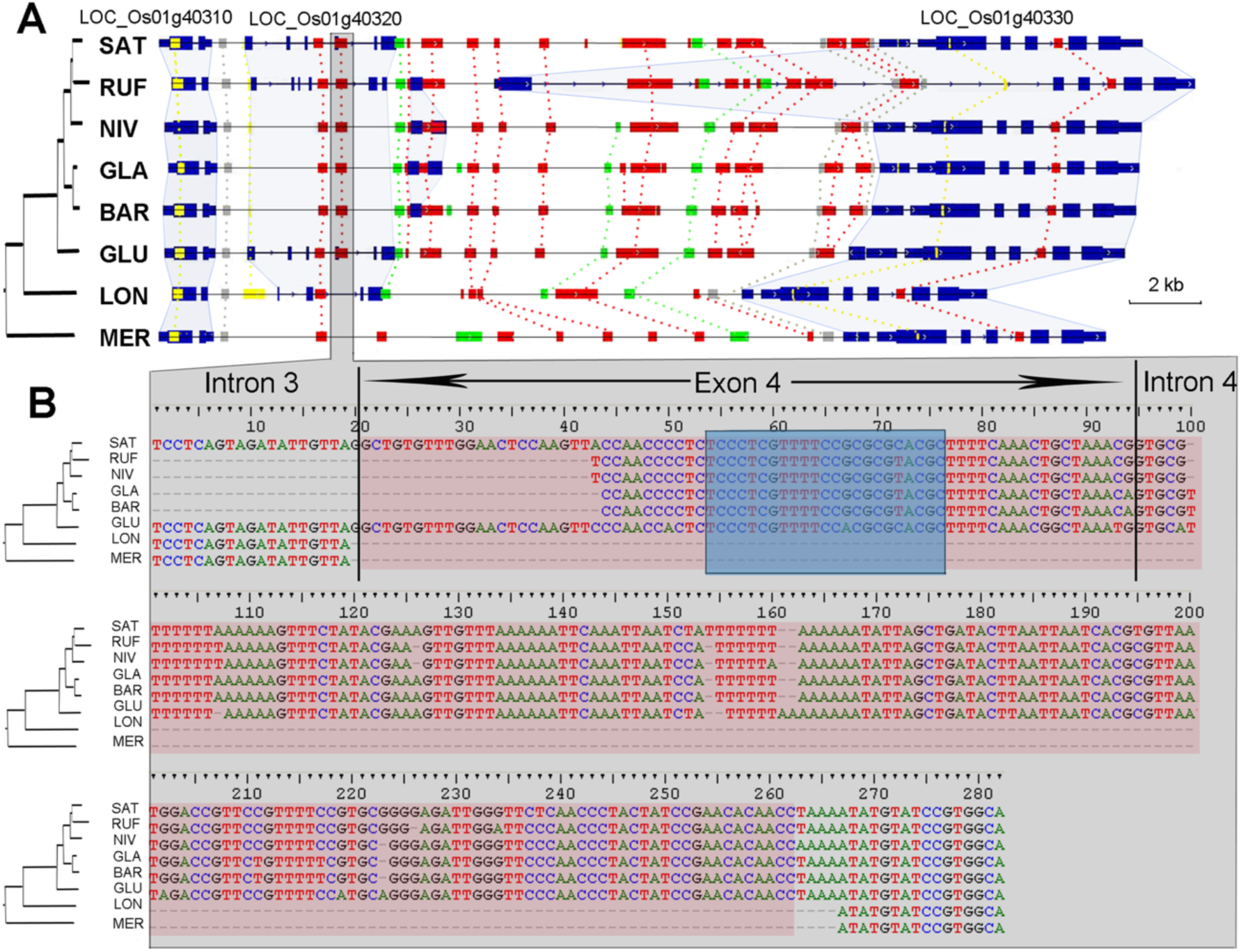
Genomic alignment of orthologous regions containing LOC_Os01g40320 across AA-genome *Oryza* species. **(A)** Orthologous genomic segments surrounding LOC_Os01g40320 in AA-genome *Oryza* species. Gene models are represented by blue rectangles. Transposable elements (TEs) are color-coded: red for DNA transposons, green for retrotransposons, and gray for unclassified types. Simple sequence repeats are shown in yellow. Dashed lines connect shared TEs; light-blue lines connect orthologous genes. **(B)** Sequence alignment of the inserted *Tourist*-like DNA transposon. The region marked with a red rectangle corresponds to the TE insertion; the region in the blue rectangle indicates the target site of osa-miR1868.

To estimate the timing of TE insertions, we constructed consensus sequences as proxies for ancestral TE families and calculated sequence divergence (mismatches relative to consensus/total sites compared) to infer insertion age. The divergence of TEs overlapping MIRNAs was significantly higher than that of TEs overlapping miRNA target sites (*P* = 0.014, t-test; Supplemental Table 6), indicating that TEs overlapping miRNA target sites are, on average, younger than those overlapping MIRNAs. Thus, TE insertions overlapping target sites likely occurred more recently and created novel miRNA target sites *de novo*.

Recognizing that our stringent MIRNA annotation criteria might have missed some young MIRNAs, we repeated the analysis using the miRBase v21 dataset. This identified 55 MIRNAs whose sequences and at least one target site overlapped the same TE. Manual inspection via the synteny map revealed further cases, such as osa-miR5825, which is associated with environmental stress response (Jeong et al., 2011). Among the 592 SAT MIRNAs in miRBase v21, osa-miR5825 is a high-confidence annotation. It is conserved across nearly all eight AA-genome species (Supplemental Figure 4) and overlaps a DNA/TcMar-Stowaway MITE. In SAT, RUF, NIV, GLA, BAR, and GLU, its target site in the 3′ UTR of LOC_Os07g37680 overlaps the same TE, whereas no TE insertion is observed at this locus in LON or MER (Supplemental Figure 5). Again, TEs overlapping this MIRNA showed significantly higher sequence divergence than those overlapping its target site (*P* < 0.001, t-test; Supplemental Table 7), confirming that the target-site-overlapping TEs are younger. Together, these results demonstrate that novel miRNA target sites can be generated *de novo* through TE insertions.

### Distinctive Evolutionary Behaviors of MiRNA Target Genes

To elucidate general principles governing the evolutionary processes of miRNA target genes—including those potentially newly generated by TE insertions—we analyzed a suite of characteristic features reflective of their evolutionary behavior. We found that miRNA target genes were significantly longer in sequence than the genomic background of all genes (*P* < 0.001, permutation test). In contrast, TE-inserted target genes (i.e., those with miRNA target sites overlapping TEs) were significantly shorter than other miRNA target genes (*P* = 0.018, permutation test; Figure 3A). The GC content of miRNA target genes was significantly lower than that of all genes (*P* < 0.001), whereas TE-inserted target genes had significantly higher GC content than other target genes (*P* < 0.001; Figure 3B).

**Figure 3.**
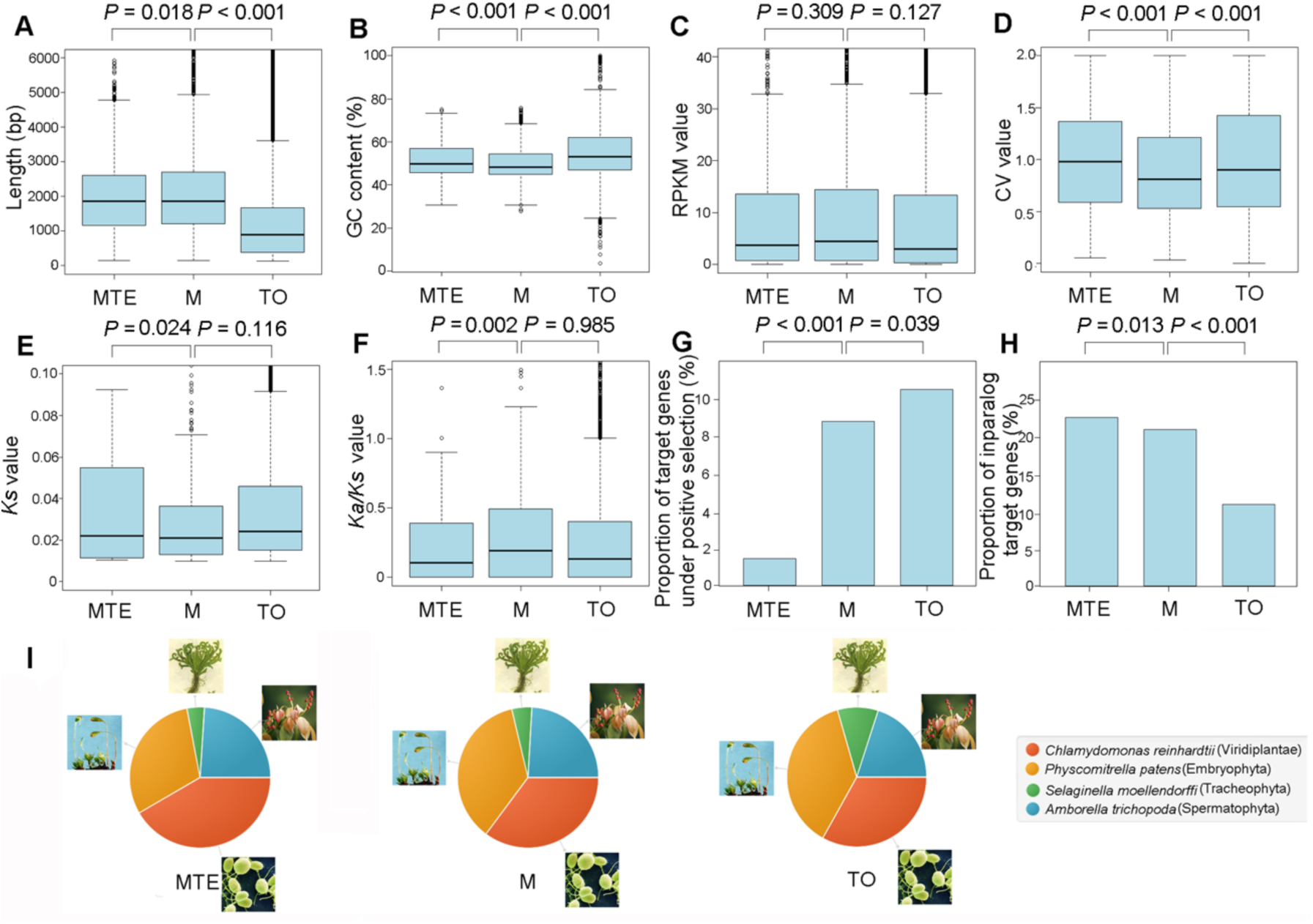
Comparison of gene features between miRNA target genes and total genes. **(A)** Gene length; **(B)** GC content; **(C)** Gene expression level; **(D)** Gene expression variability; **(E)** *Ks* value; **(F)** *Ka*/*Ks* ratio; **(G)** Proportion of target genes under positive selection; **(H)** Proportion of inparalog target genes; **(I)** Distribution of evolutionary ages. MTE, miRNA target genes with TE-overlapping target sites; M, all miRNA target genes; TO, total genes. *P*-values in (A)–(F) were derived from permutation tests; *P*-values in (G) and (H) were obtained by chi-square tests.

We next assessed expression patterns by calculating RPKM (Reads Per Kilobase of transcript per Million mapped reads, expression level) and the coefficient of variation (CV, expression variability). miRNA target genes showed moderately higher expression levels than all genes, though this difference was not significant (*P* = 0.127). The expression levels of TE-inserted target genes did not differ significantly from other target genes (*P* = 0.309; Figure 3C). However, miRNA target genes exhibited significantly lower expression variability than all genes (*P* < 0.001), while TE-inserted target genes showed significantly higher variability than other target genes (*P* < 0.001; Figure 3D).

To quantify nucleotide substitution and selective pressure, we estimated the synonymous substitution rate (*Ks*) and the *Ka*/*Ks* ratio using a branch model in PAML (Yang, 2007), and identified genes under positive selection using a branch-site model. Compared to all genes, miRNA target genes had relatively low nucleotide substitution rates (*P* = 0.116), non-significantly different *Ka*/*Ks* values (*P* = 0.985), and a significantly lower proportion of genes under positive selection (*P* = 0.039; Figure 3E–G). Relative to other miRNA target genes, TE-inserted target genes had significantly higher nucleotide substitution rates (*P* = 0.024), lower *Ka*/*Ks* values (*P* = 0.002), and a lower proportion of genes under positive selection (*P* < 0.001; Figure 3E–G).

We also examined recent gene duplication using orthoMCL (Fischer et al., 2011) to identify inparalogs. miRNA target genes, especially TE-inserted ones, were significantly more likely to be inparalogs compared to all genes (*P* < 0.001; Figure 3H).

Gene ages were estimated via phylostratigraphy (Arendsee et al., 2014), categorizing genes into four evolutionary stages: originating before Viridiplantae (∼968 Myr), Embryophyta (∼547 Myr), Tracheophyta (∼496 Myr), or Spermatophyta (∼329 Myr). A significantly higher proportion of miRNA target genes originated in the earliest stage (before Viridiplantae) compared to all genes (35.16% vs. 33.08%, *P* = 0.009, chi-square test; Figure 3I). This trend was even more pronounced for TE-inserted target genes, which showed a higher early-origin proportion than other miRNA target genes (41.60% vs. 35.16%, *P* < 0.001; Figure 3I). Gene Ontology enrichment analysis revealed that TE-inserted target genes were significantly associated with secondary metabolic processes (GO:0019748; Supplemental Figure 6). Likewise, a consistent pattern was observed for the subset of miRNA target genes where both the MIRNA and its target site overlapped with the same TE (Supplemental Figure 7).

In summary, miRNA target genes possess distinct evolutionary features compared to the genomic background, and TE-inserted target genes further exhibit distinguishable characteristics within the miRNA target set, highlighting their unique evolutionary trajectories.

### Coupled evolution of MIRNAs with target genes and secondary silencing pathways

To elucidate the potential co-evolutionary interplay between MIRNAs and their target genes, we analyzed the features of target genes regulated by different MIRNA groups: duplication-generated, *de novo* generated, and conserved MIRNAs. Duplication-generated MIRNAs are newly evolved loci that have homologs present at the ancestral node of the AA-genome *Oryza* phylogeny (22.67% overlap with TEs). *De novo* generated MIRNAs are also newly evolved but lack homologs at that ancestral node (41.18% overlap with TEs; *P* < 0.001 compared with duplication-generated MIRNAs). Conserved MIRNAs are those already present at the ancestral node (16.93% overlap with TEs; *P* = 0.013 compared with duplication-generated, *P* < 0.001 compared with *de novo*generated MIRNAs).

Compared with target genes of *de novo* generated and conserved MIRNAs, target genes of duplication-generated MIRNAs were significantly longer (*P* < 0.001, permutation test; Figure 4A) and had lower GC content (*P* < 0.001; Figure 4B)—a pattern also observed in the corresponding MIRNA and mature miRNA sequences (Supplemental Figure 8). Their expression levels and variability did not differ significantly from other groups (*P* = 0.278 and *P* = 0.763, respectively; Figure 4C, D). However, they showed higher nucleotide substitution rates (*P* = 0.043), marginally higher *Ka*/*Ks* values (*P* = 0.073), and a similar proportion of genes under positive selection (*P* = 0.500) compared to other target genes (Figure 4E–G). They were also more likely to be inparalogs (*P* < 0.001; Figure 4H).

**Figure 4.**
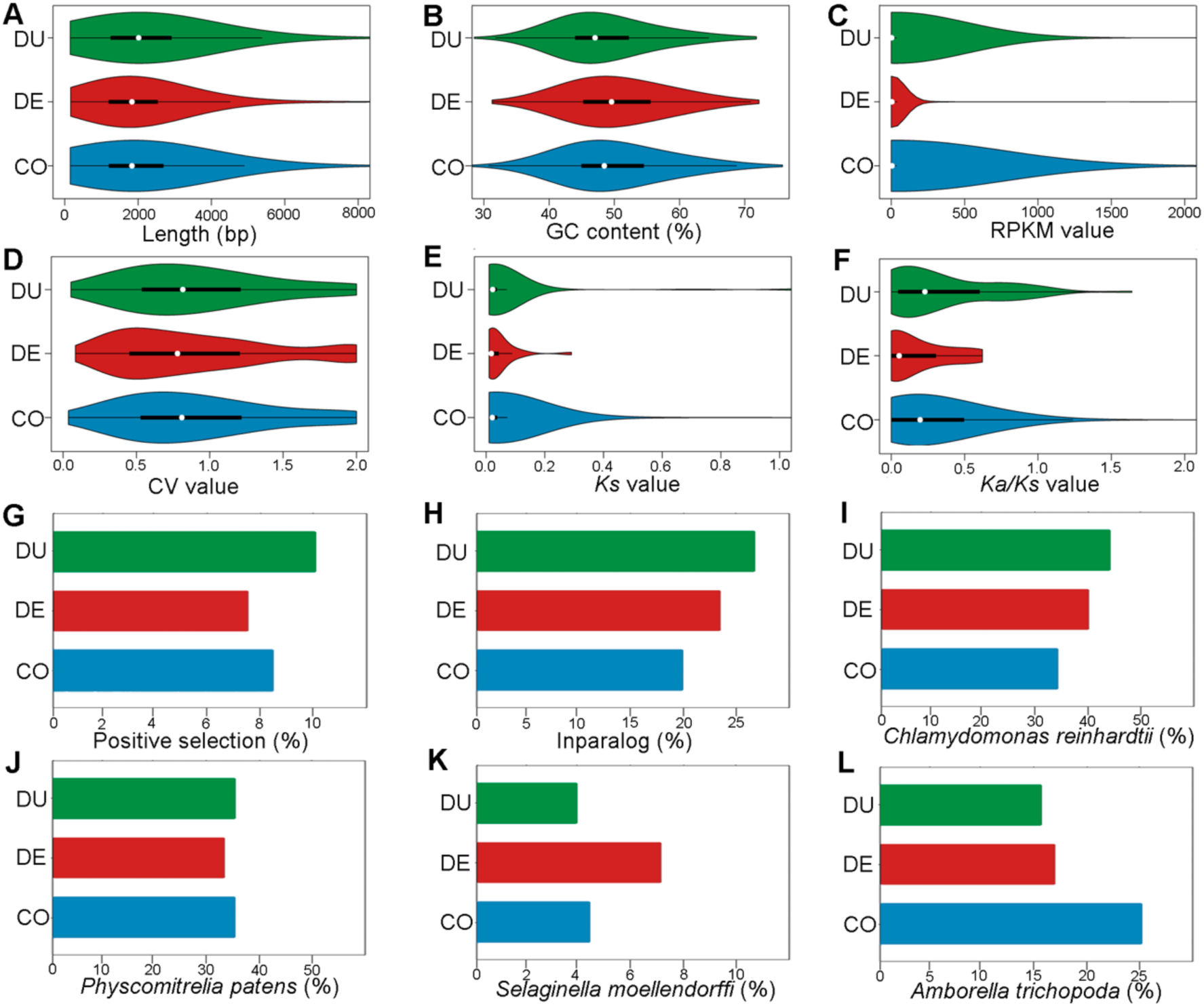
Comparison of target-gene features among *de novo* generated, duplication-generated, and conserved MIRNAs. **(A)** Gene length; **(B)** GC content; **(C)** Gene expression level; **(D)** Gene expression variability; (E) *Ks* value; (F) *Ka*/*Ks* ratio; **(G)** Proportion of target genes under positive selection; **(H)** Proportion of inparalog target genes; **(I)** Proportion of target genes originating before Viridiplantae (*Chlamydomonas reinhardtii*); **(J)** Proportion originating before Embryophyta (*Physcomitrella patens*); **(K)** Proportion originating before Tracheophyta (*Selaginella moellendorffii*); **(L)** Proportion originating before Spermatophyta (*Amborella trichopoda*). DU, targets of duplication-generated MIRNAs; DE, targets of *de novo* generated MIRNAs; CO, targets of conserved MIRNAs.

Interestingly, newly generated MIRNAs—especially duplication-generated ones—showed a bias toward regulating evolutionarily old genes rather than young ones. Compared with targets of conserved MIRNAs, targets of *de novo* generated MIRNAs contained a higher proportion of ancient genes (originating before Viridiplantae: 40.91% vs. 34.00%; *P* = 0.160) and a lower proportion of young genes (originating before Spermatophyta: 17.27% vs. 25.25%; *P* = 0.073) (Figure 4I, L). For duplication-generated MIRNAs, this trend was stronger and statistically significant: their target genes had a markedly higher proportion of ancient genes (44.26% vs. 34.00%; *P* < 0.001) and a lower proportion of young genes (15.55% vs. 25.25%; *P* < 0.001) (Figure 4I, L). Gene Ontology enrichment analysis revealed no significant terms for targets of *de novo* generated MIRNAs, whereas targets of duplication-generated MIRNAs were enriched for developmental process (GO:0032502) and multicellular organismal process (GO:0032501) (Supplemental Figure 9). Together, these results indicate distinct evolutionary profiles among target genes of different MIRNA origins, with duplication-generated MIRNAs in particular regulating an older, development-related gene set, suggesting group-specific co-evolutionary dynamics.

To further examine co-evolution between MIRNAs and secondary silencing pathways, we analyzed phased siRNAs (phasiRNAs) produced from miRNA target genes. Using ShortStack (Shahid and Axtell, 2014), we identified 2,719 *PHAS* loci overlapping miRNA target genes across the eight AA-genome species. Phasing scores were calculated as described by Guo and colleagues (Guo et al., 2015), with higher scores indicating stronger phasing signals. We found that *PHAS* loci associated with targets of *de novo* generated MIRNAs were longer (*P* < 0.001; Figure 5A), had lower siRNA abundance (*P* = 0.016; Figure 5B), and exhibited lower phasing scores (*P* = 0.040; Figure 5C) than those of other target genes. However, a higher proportion of target genes of *de novo* generated MIRNAs were associated with phasiRNA-producing *PHAS* loci (*P* < 0.001, chi-square test; Figure 5D). These findings suggest that *PHAS* loci co-evolving with MIRNAs have undergone distinct evolutionary trajectories, which may reflect previously unrecognized steps in the maturation of miRNA–phasiRNA regulatory modules.

**Figure 5.**
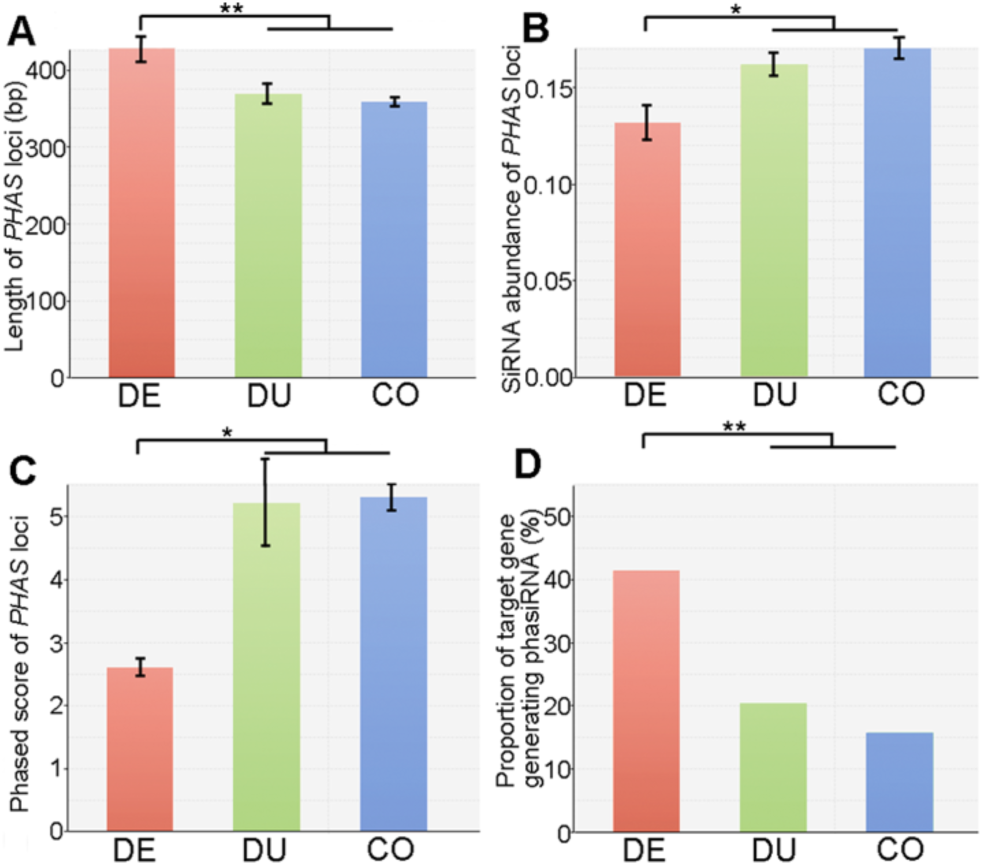
Comparison of *PHAS* locus features among target genes of *de novo* generated, duplication-generated, and conserved MIRNAs. **(A)** Length of *PHAS* loci; **(B)** siRNA abundance of *PHAS* loci; **(C)** Phasing score of *PHAS*loci; **(D)** Proportion of target genes producing phasiRNA. DE, target genes of *de novo* generated MIRNAs; DU, target genes of duplication-generated MIRNAs; CO, target genes of conserved MIRNAs. Data in **(A)**–**(C)** are presented as mean ± s.e.m.; *P*-values were derived from permutation tests. *P*-value in **(D)** was obtained by chi-square test. Asterisks denote statistical significance: \*\**P* < 0.001, \**P* < 0.05.

## DISCUSSION

To understand MIRNA evolution in plants, accurate genome-wide annotation is essential yet challenging. The number of MIRNAs we annotated is lower than in some previous reports (Kozomara and Griffiths-Jones, 2014). Although we may have missed some genuine loci, much of this discrepancy likely stems from the inclusion of incorrectly annotated entries in public databases that do not meet strict MIRNA criteria (Meng et al., 2012; Taylor et al., 2014). Such annotation issues have contributed to the “rapid turnover” model, which posits a high birth rate of new MIRNA families that are often quickly lost (Fahlgren et al., 2007). However, the apparent abundance of species-specific families may reflect accumulated annotation errors and sparse phylogenetic sampling rather than true evolutionary dynamics (Taylor et al., 2014). Our phylogenetically informed analysis reconciles these views: while many hairpin-derived RNAs may be evolutionarily transient, bona fide MIRNA families actually exhibit low birth–death rates, whereas individual paralogous MIRNAs turn over rapidly.

The inverted-duplication model explains some *de novo* MIRNA origins from protein-coding sequences (Allen et al., 2004), yet it cannot readily account for MIRNAs that regulate genes with little sequence similarity (Takuno and Innan, 2008). Notably, some TEs themselves can be targeted by miRNAs, leading to epigenetically activated siRNAs (Creasey, Zhai et al., 2014). If a MIRNA arises from one gene family via inverted duplication, it remains unclear how target sites could emerge in unrelated gene families or in TEs. Strongly supporting our earlier hypothesis (Zhang et al., 2011), we demonstrate here that novel miRNA target sites can be generated *de novo* through TE insertions into protein-coding genes, as shown across all eight AA-genome *Oryza* species. The target sites appear to have been inserted after the MIRNAs themselves originated. We therefore propose a “TE-MIRNA co-option” model: one TE copy evolves into a MIRNA, while another copy of the same TE family inserts into a protein-coding gene to create a new miRNA target site. If validated, this mechanism could offer a novel strategy for engineering miRNA-mediated regulatory circuits to improve crop traits, obviating the need for introducing exogenous MIRNAs (Chen et al., 2016; Zhao et al., 2015).

miRNA target genes display distinct evolutionary features compared with typical protein-coding genes. They tend to be longer, lower in GC content, moderately highly expressed, with low expression variability, relatively low nucleotide-substitution rates, and weaker selective pressures. Genes with such properties are often ancient, conserved, and slow-evolving, in contrast to faster-evolving genes that are influenced by processes such as GC-biased gene conversion (Carels and Bernardi, 2000; Muyle et al., 2011; Ren et al., 2006; Serres-Giardi et al., 2012; Tatarinova et al., 2010; Vishnoi et al., 2010).

Intriguingly, despite the repressive role of miRNAs, their target genes exhibited a non-significant trend toward higher expression compared to the genomic background. This observation suggests that miRNA regulation may be preferentially recruited to tune the expression of inherently active genes within the conserved “kernel” subsystem. The actual expression potential of these targets might be even greater than observed, as miRNA-mediated suppression likely modulates their levels to maintain optimal expression ranges and buffer against perturbations.

Our phylostratigraphic dating confirms that miRNA targets are enriched for early-originating genes. In animal systems, gene-regulatory networks are thought to comprise a conserved, slowly evolving “kernel” subsystem alongside more rapidly evolving “plug-in” components (Chen and Rajewsky, 2007). Although plants have fewer miRNA-regulated genes than animals, our findings suggest that miRNAs preferentially enhance the tunability and robustness of the plant “kernel” developmental subsystem. Within this subsystem, where gene-expression programs face both internal and external perturbations, certain core genes may gain a selective advantage by being linked to miRNAs, which can act as expression “guardians” (Leung and Sharp, 2007).

Our results indicate that a novel mechanism may exist to acquire miRNA target sites through TE insertions. Our results further reveal that TE-inserted target genes possess a distinct evolutionary profile: they are shorter (despite the added TE sequence), higher in GC content, show higher expression variability, higher nucleotide-substitution rates, lower *Ka*/*Ks* values, a lower proportion under positive selection, and a higher proportion of ancient genes. These traits collectively indicate a relatively accelerated evolutionary pace.

The association between newly evolved miRNAs and TE-embedded targets raises the question of its functional significance. Interestingly, newly generated 24-nt MIRNAs have been shown to direct DNA methylation of their targets (Wu et al., 2010), and we previously observed higher methylation levels in young *versus* conserved MIRNAs in *Arabidopsis* (Zhang et al., 2011). Thus, RNA-directed DNA methylation might serve as a pathway to suppress TE activity. We speculate that newly evolved miRNAs, by partnering with TE-located target genes, could help control TE mobility, adding a layer of functional constraint to this apparently fast-evolving regulatory module.

Recent studies indicate that newly generated MIRNAs often regulate secondary-metabolite synthesis (Li et al., 2015; Sharma et al., 2016). Consistently, our GO analysis shows enrichment of secondary-metabolic processes among TE-inserted target genes. A plausible interpretation is that genes involved in secondary metabolism—which function in defense or specific developmental programs (Fei et al., 2013; Xia et al., 2013)—are preferentially regulated by newly evolved MIRNAs and belong to the more rapidly evolving fraction of the “kernel” subsystem.

Although MIRNA–target co-evolution has long been hypothesized (Tang et al., 2010; Zhang et al., 2011), empirical evidence in plants remains scarce. We found that MIRNAs, particularly those generated by duplication, preferentially target inparalogs produced by recent gene duplications. Despite being ancient (long, low-GC, early-originating), targets of duplication-generated MIRNAs show elevated nucleotide-substitution rates and slightly higher *Ka*/*Ks* values, indicating moderate evolutionary change. What drives this distinct pattern? Increased complexity in multicellular development often arises from duplication and subsequent divergence of ancient genes, coupled with regulatory diversification, rather than from entirely novel genes (Conant and Wolfe, 2008). Our GO analysis reveals that targets of duplication-generated MIRNAs are enriched in developmental and multicellular-organismal processes, and MIRNA duplications themselves are frequent across the AA-genome lineages. Thus, a coherent explanation emerges: as the “kernel” developmental subsystem gains complexity through gene duplication, the corresponding MIRNA regulators also duplicate and evolve, thereby fine-tuning and buffering the expanding network. Whereas the gain of entirely new MIRNA families may not directly correlate with phenotypic change in plants as it does in animals (Taylor et al., 2014), the duplication and subsequent evolution of existing MIRNAs could represent an equivalent route to increasing regulatory complexity within the plant miRNAome.

Finally, miRNAs can trigger phased siRNAs (phasiRNAs) to reinforce silencing, sometimes acting as non-cell-autonomous signals. Although these secondary siRNA pathways are ancient, their evolutionary mechanisms remain poorly understood (Fei et al., 2013). We observed that *PHAS* loci associated with targets of *de novo* generated MIRNAs are longer, produce lower siRNA abundance, have lower phasing scores, yet are more frequently associated with phasiRNA production. This suggests that as *de novo* MIRNAs mature, their linked *PHAS* loci also evolve toward more focused and efficient phasiRNA output at fewer loci. Thereby, the co-evolution of secondary siRNAs adds another layer of tunability and robustness to the regulatory system.

## METHODS

### Small RNA Sequencing and Data Processing

Seven AA-genome *Oryza* accessions—*O. rufipogon* (RUF), *O. nivara* (NIV), *O. glaberrima* (GLA), *O. barthii* (BAR), *O. glumaepatula* (GLU), *O. longistaminata* (LON), and *O. meridionalis* (MER)—were obtained from the International Rice Research Institute (IRRI) for small RNA sequencing. Roots and shoots from one-month-old seedlings, as well as flag leaves and panicles at the booting stage from greenhouse-grown plants, were collected, immediately frozen in liquid nitrogen, and stored at −80 °C until RNA extraction. Total RNA was isolated using a water-saturated phenol method (Zhang et al., 2014). Approximately 30 μg of total RNA was separated on a 15% acrylamide/8 M urea gel, and small RNAs corresponding to 19- and 24-nt fragments were excised and recovered using a high-salt extraction protocol. Recovered small RNAs were ligated to a 3′ adaptor (IDT), and the ligated products were size-selected on a 15% acrylamide/8 M urea gel. Gel fragments containing 37–42 nt molecules were excised, and the RNA was purified. A 5′ sequencing adaptor was added with T4 RNA ligase, followed by reverse transcription using Superscript III (Invitrogen) and PCR amplification. The resulting cDNA libraries were size-selected on 3% agarose gels; bands corresponding to 108–115 nt were excised, purified, and sequenced on an Illumina platform.

Publicly available genome and transcriptome data for RUF, NIV, GLA, BAR, GLU, LON and MER were obtained from http://www.plantkingdomgdb.com. The genome and transcriptome of *O. sativa* ssp. *japonica* cv. Nipponbare (SAT) were downloaded from the Rice Genome Annotation Project (release 7.0; http://rice.plantbiology.msu.edu). Small RNA data for SAT were acquired from the Cereal Small RNA Database (http://sundarlab.ucdavis.edu/smrnas/). Genomes of *O. brachyantha* and *Sorghum bicolor* were retrieved from Ensembl Plants (Bolser et al., 2016). Prior to analysis, all TE-related and non-coding RNA genes were excluded. For genes with alternative splicing isoforms, the longest transcript was selected.

Raw small-RNA reads were filtered to remove low-quality sequences, and adapter trimming, read collapsing, and size selection (19–24 nt) were performed with the fastx-toolkit (http://hannonlab.cshl.edu/fastx_toolkit/). *De novo* identification of MIRNAs in each *Oryza* genome was conducted using the miRDeep-P pipeline (Yang and Li, 2011). Briefly, small-RNA libraries from the four tissues of each species were concatenated, mapped to the respective genome, and candidate hairpin structures were predicted using RNA folding. Processing precision and the presence of a miRNA* sequence were evaluated for each precursor. Loci lacking a clear dominant mature product were excluded; imprecisely processed loci were retained only when a homologous locus meeting annotation criteria was found in another *Oryza* lineage. Conserved MIRNAs were identified by BLAST searches against miRBase v21. Iterative homology searches were performed between each miRDeep-P-derived miRNAome and each genome, with careful validation of potential homologs. A final list of MIRNAs per genome was compiled and grouped into families based on mature-sequence similarity. Homology searches were extended to the outgroup genomes of *S. bicolor* and *O. brachyantha*.

### Annotation of Transposable Elements

Previously annotated rice transposons were downloaded from Repbase (http://www.girinst.org/repbase/). *De novo* repeat families were identified and modeled using RepeatModeler (http://www.repeatmasker.org/RepeatModeler.html), which incorporates RECON and RepeatScout. Combined with Repbase DNA-transposon entries, the RepeatModeler output was used to build a custom DNA-transposon library for RepeatMasker (Tempel, 2012) to annotate TEs in the eight AA-genome *Oryza* assemblies. LTR retrotransposons were identified *de novo* using LTR_STRUCT (McCarthy and McDonald, 2003), and predictions were verified with LTR_FINDER (Xu and Wang, 2007). Intact LTR retrotransposons were classified into families using BLASTClust, applying the criterion that 5′ LTR sequences of the same family share ≥80% identity over ≥80% of their length (Seberg and Petersen, 2009). PFAM (http://pfam.sanger.ac.uk) was run against the intact LTR retrotransposon library to determine protein-domain order. Retrotransposons were classified into *Ty3-gypsy* (PR-RT-RH-IN), *Ty1-copia*(PR-IN-RT-RH), or unclassified superfamilies based on the arrangement of reverse-transcriptase (RT), integrase (IN), protease (PR) and RNaseH (RH) domains in the *pol* gene (Coffin JM, 1997). LTR retrotransposons encoding RT were further clustered into families using BLASTClust with a threshold of >80% identity over >80% of the RT protein length. Finally, the custom *Oryza* TE library and Repbase LTR retrotransposon annotations were merged to create a comprehensive RepeatMasker library for annotating LTR retrotransposons in the eight AA-genomes.

### Construction of the Orthologous Synteny Map of Eight *Oryza* AA-genomes

Orthologous genomic regions across the eight AA-genome *Oryza* species were identified and aligned using MERCATOR and MAVID(Bray and Pachter, 2004). The procedure consisted of six steps: (1) all exons of a reference genome were aligned to exons of the other seven genomes using BLAT, and significant exon-level alignments were recorded; (2) a graph was built with vertices representing exons and edges connecting exons with significant alignments; (3) cliques in the graph were identified; (4) adjacent cliques were merged into runs; (5) the genomic span of each run was output as an orthologous segment; (6) orthologous sequences were aligned under the established phylogenetic relationships: ((((((SAT:0.002022, RUF:0.003527):0.000373, NIV:0.002659):0.001616, (BAR:0.001242, GLA:0.001793):0.001899):0.001268, GLU:0.004631):0.007018, LON:0.011113):0.003953, MER:0.014768) (Zhu et al., 2014). In total, 8,742 orthologous genomic segments were obtained (Supplemental Table 8).

### Analysis of MiRNA Target Genes

miRNA target genes were predicted separately for each AA-genome species using psRNATarget (Dai and Zhao, 2011)with default parameters: maximum expectation 3.0, complementary scoring length 20 bp, maximum unpairing energy for target accessibility 25.0, flanking regions of 17 bp upstream and 13 bp downstream of the target site, and central mismatch region of 9–11 bp for translational inhibition. Gene expression levels (RPKM) were calculated with RSEM (Haas et al., 2013) using the genome as reference; the mean RPKM across tissues was used as the expression level for each gene. Expression variability was defined as the coefficient of variation (CV; standard deviation divided by the mean).

To assess duplication status and selection pressures, orthoMCL (Fischer et al., 2011) was first used to identify inparalogs and orthologs within each species via an all-vs-all BLASTP search (e-value < 10⁻⁵) and Markov clustering. High-confidence orthologous gene sets were obtained by intersecting orthoMCL results with orthologous segments defined by MERCATOR, followed by alignment with ClustalW. Alignments containing frameshift indels, CDS lengths not a multiple of three, in-frame stop codons, or sequences outside MERCATOR orthologous segments were discarded, yielding 6,501 orthologous gene sets for downstream analysis. Protein alignments were converted to codon alignments using Bioperl scripts.

The “two-ratio” branch model in codeml (PAML) (Yang, 2007)was used to estimate branch-specific *Ks* and *Ka*/*Ks* values. Genes with *Ks* between 0.01 and 2 were retained (Yang, 1998). Positive selection was tested using a branch-site model: the alternative model (positive selection) was compared with the null model (nearly neutral, ω₂ = 1 fixed) via a likelihood-ratio test (LRT). Sites under positive selection were identified with the Bayes empirical Bayes (BEB) method. *P*-values were computed assuming a 50:50 mixture of a point mass at 0 and χ² with 1 d.f., then adjusted for false-discovery rate (FDR) using the Benjamini–Hochberg procedure (Anisimova and Yang, 2007). Genes with FDR < 0.05 were considered under positive selection.

Gene ages were estimated by phylostratigraphy using orthology data from Ensembl Plants (Bolser et al., 2016) for *Chlamydomonas reinhardtii*, *Physcomitrella patens*, *Selaginella moellendorffii* and *Amborella trichopoda* relative to SAT. Only high-confidence ortholog pairs were retained. GO enrichment analysis of miRNA target genes was performed with agriGO (http://bioinfo.cau.edu.cn/agriGO/), using all SAT protein-coding genes as background. Fisher’s exact test with Yekutieli multi-test adjustment was applied (significance threshold = 0.05). For genes in the wild rice species, orthologs in SAT were used for age and GO analyses. Phased siRNA-producing loci (*PHAS* loci) were identified with ShortStack (Shahid and Axtell, 2014) under default settings. Phasing scores were calculated as described by Guo et al. (Guo et al., 2015).

### Statistical Analysis of Gene Features

Because distributions of variables such as GC content and *Ka*/*Ks* deviated from normality, permutation tests were employed to compare group means (e.g., miRNA target genes vs. all genes). Briefly, the observed mean difference (*t*) between two groups of sizes X and Y was computed. All genes were pooled, and a random sample of X genes was drawn to represent the first group, leaving Y genes for the second group; the difference (*t*⁺) was recorded. This resampling was repeated 10,000 times via Monte Carlo simulation. The *P*-value was calculated as the proportion of permutations where *t*⁺ exceeded *t*: *P* = (number of *t*⁺ > *t*) / number of *t*⁺.

## Supporting information

Supplemental Table 1. MIRNAs in the eight AA- genome Oryza species

Supplemental Table 4. Summary of MIRNA genes and miRNA target sites overlapped with the same TEs

## ACKNOWLEDGMENTS

We thank the International Rice Research Institute (Manila, Philippines) for providing the rice germplasm. This work was supported by the Yunnan Innovation Team Project, and a start-up grant from Hainan University (to L.G.).

## AUTHOR CONTRTRIBUTIONS

L.-Z.G. and P. D. designed the study. L.-P. Z. performed the experiments. Y.Z. and R.S.T. analyzed the data. L.-Z.G. and Y.Z. drafted the manuscript. L.-Z.G., P. D., and Y.Z. revised the manuscript.

## Supplemental Information

### Supplemental Figures

**Supplemental Figure 1.**
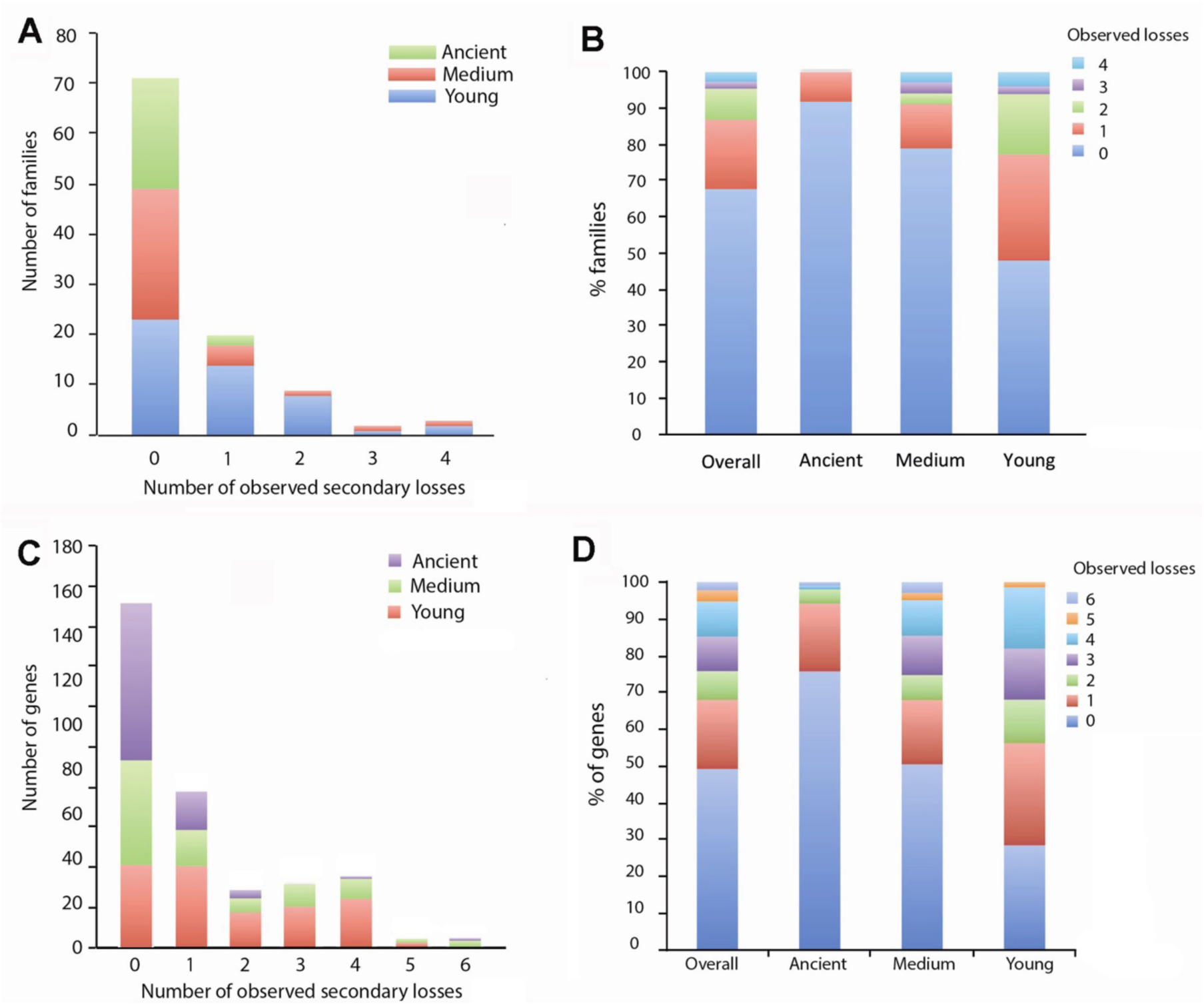
Distribution of evolutionary loss rates and ages of MIRNAs. (A, B) MIRNA families; (C, D) Individual MIRNAs.

**Supplemental Figure 2.**
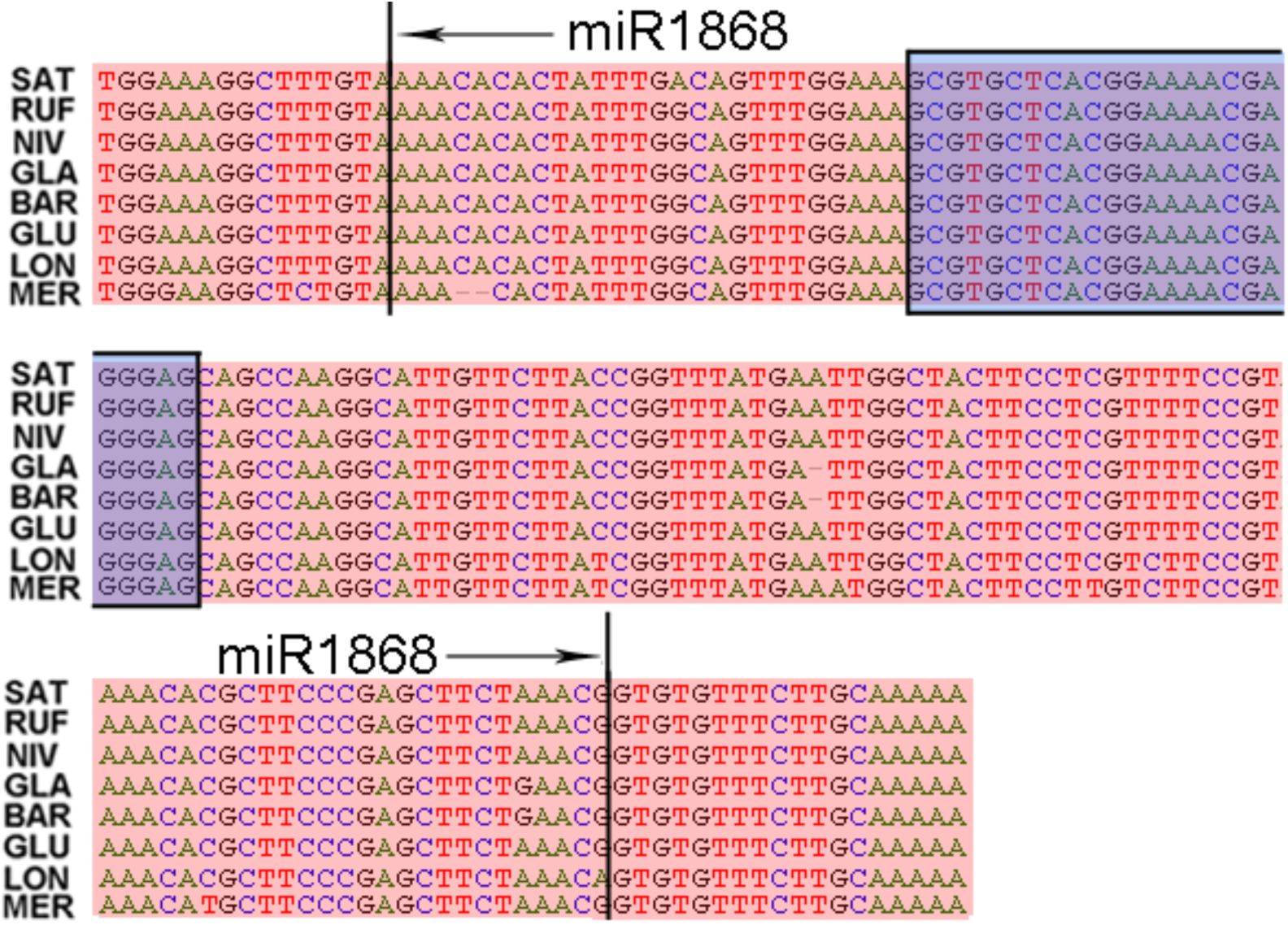
Alignment of osa-miR1868 orthologs across the eight AA-genome *Oryza* species. Sequences highlighted in the red box correspond to TEs (DNA transposon/*Tourist*); sequences in the blue box represent the mature miRNA within the MIRNAs.

**Supplemental Figure 3.**
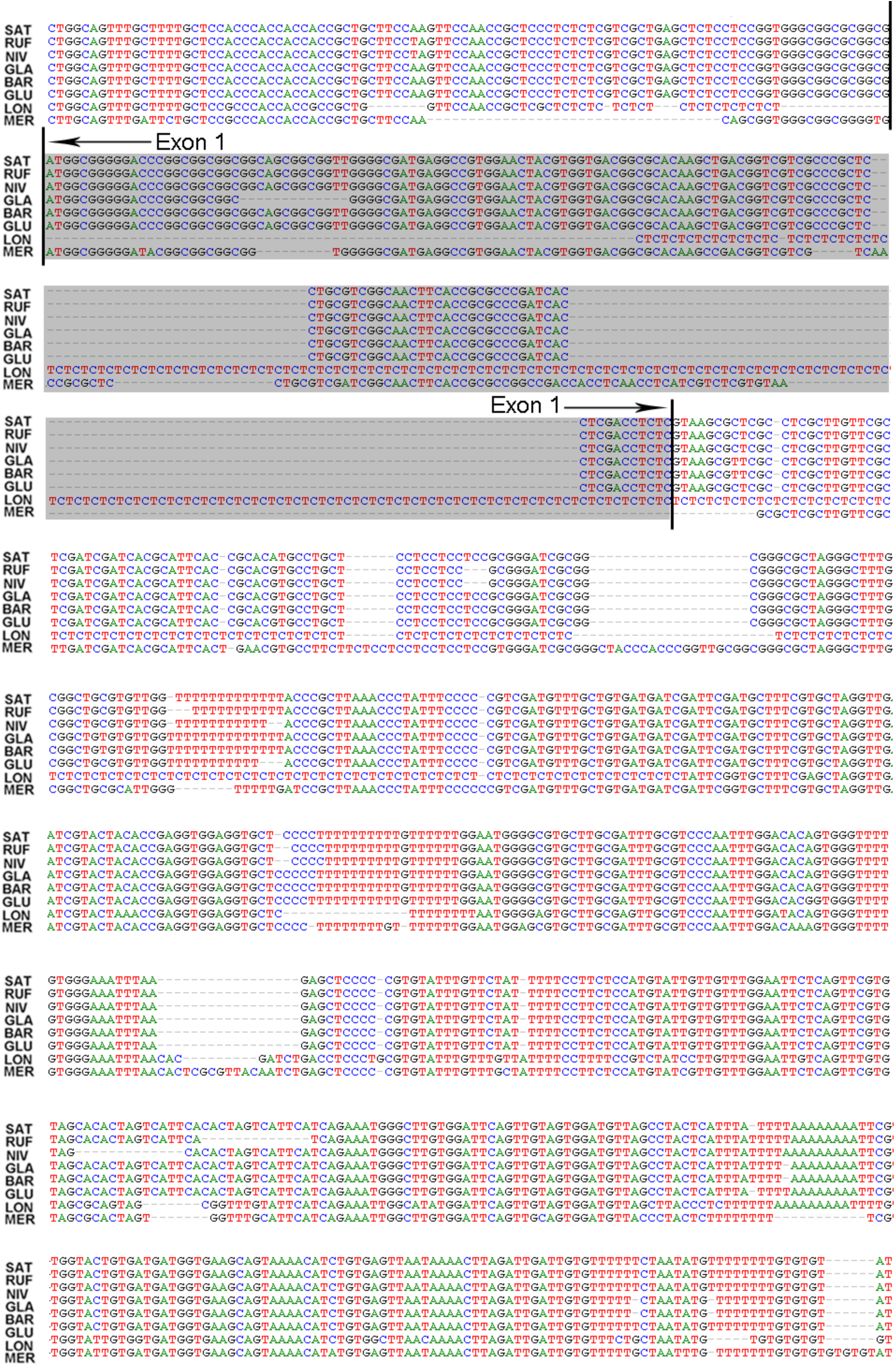
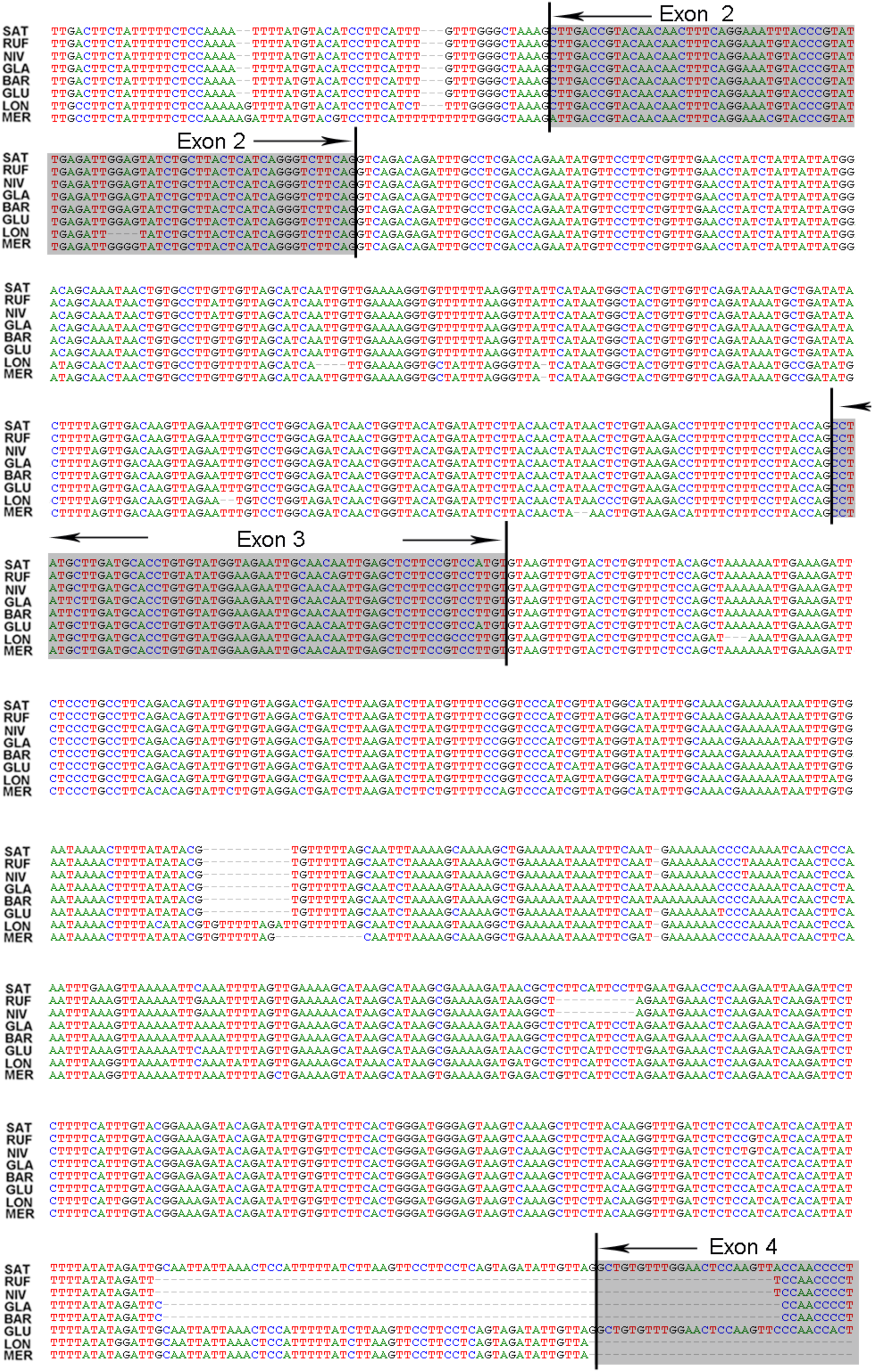
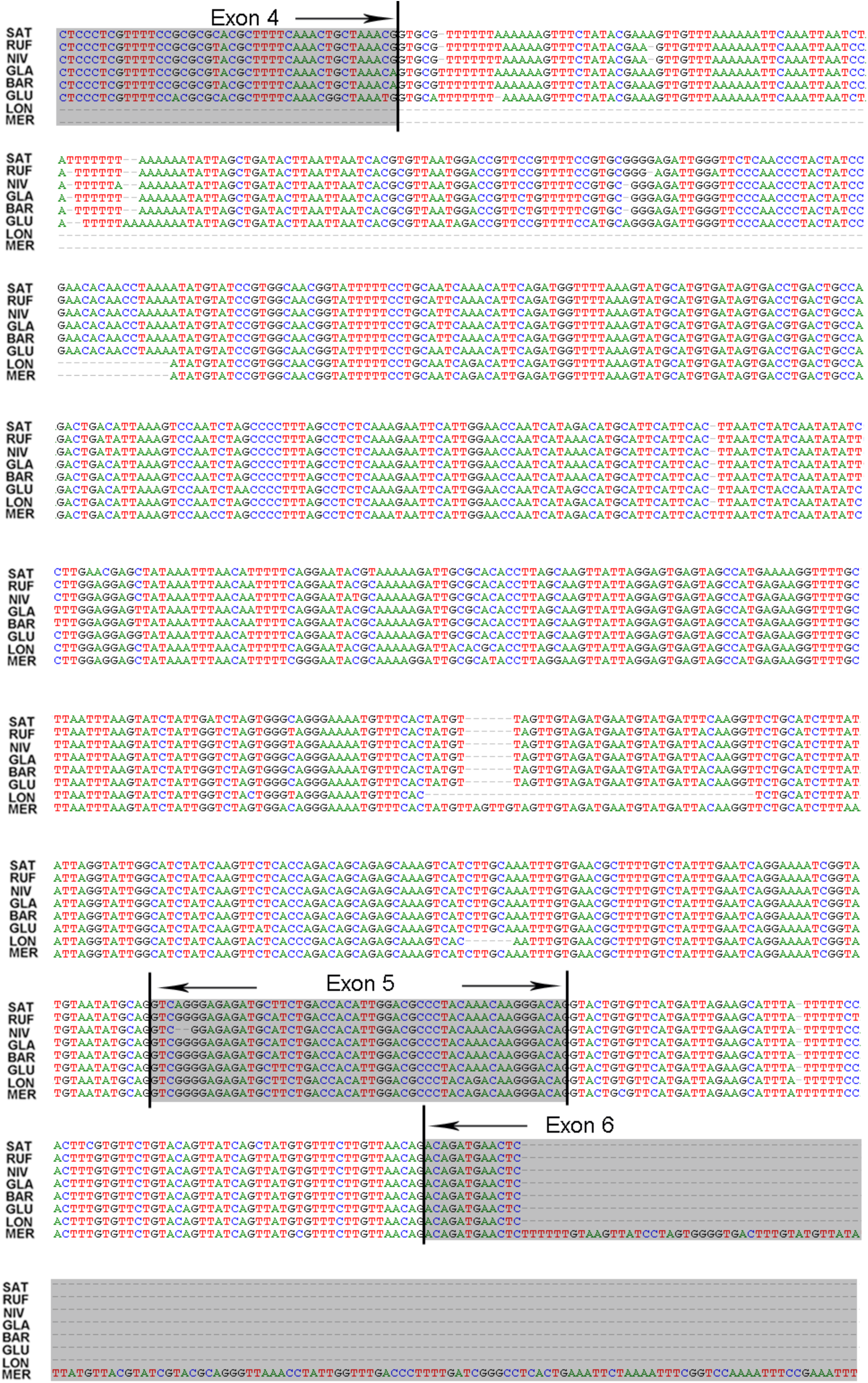
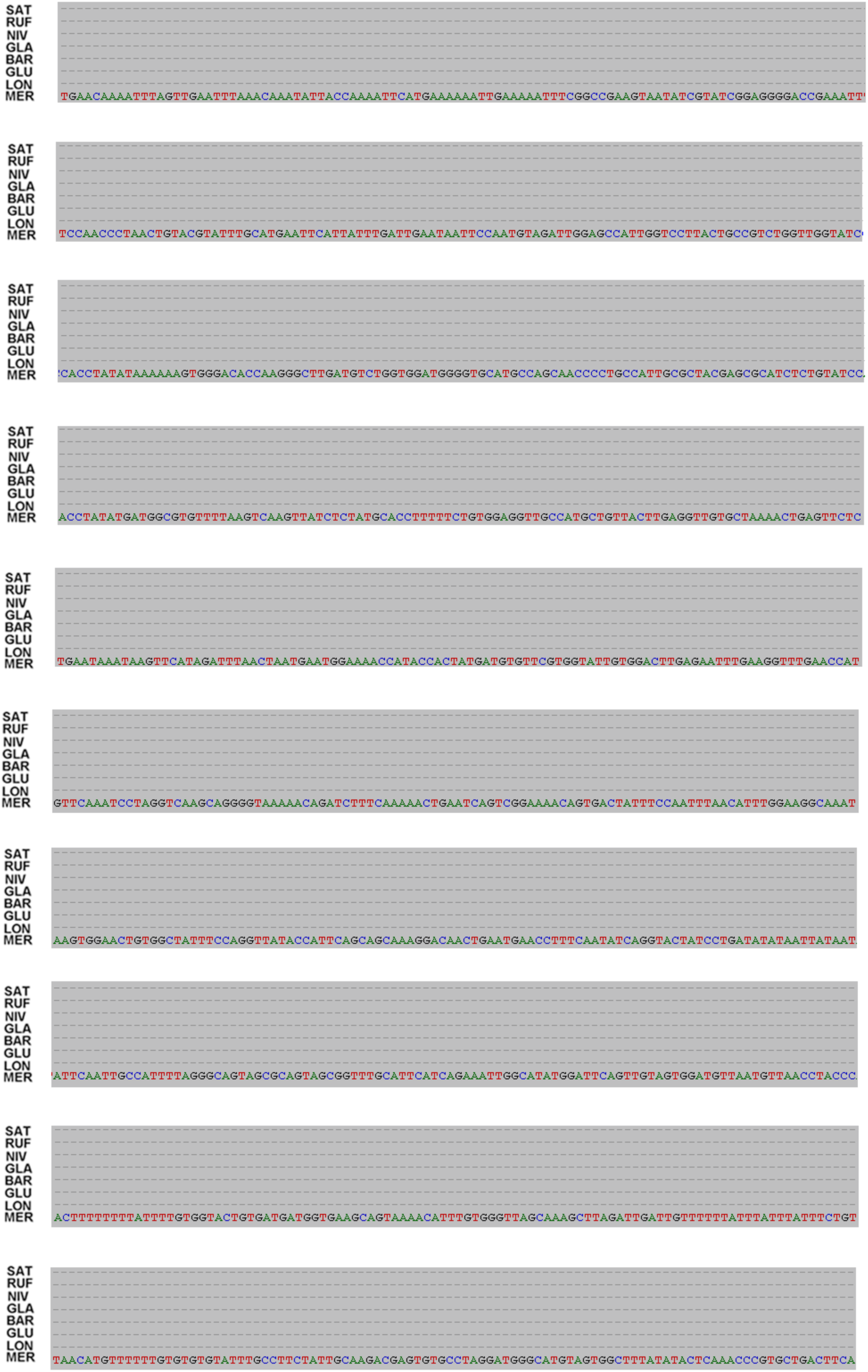
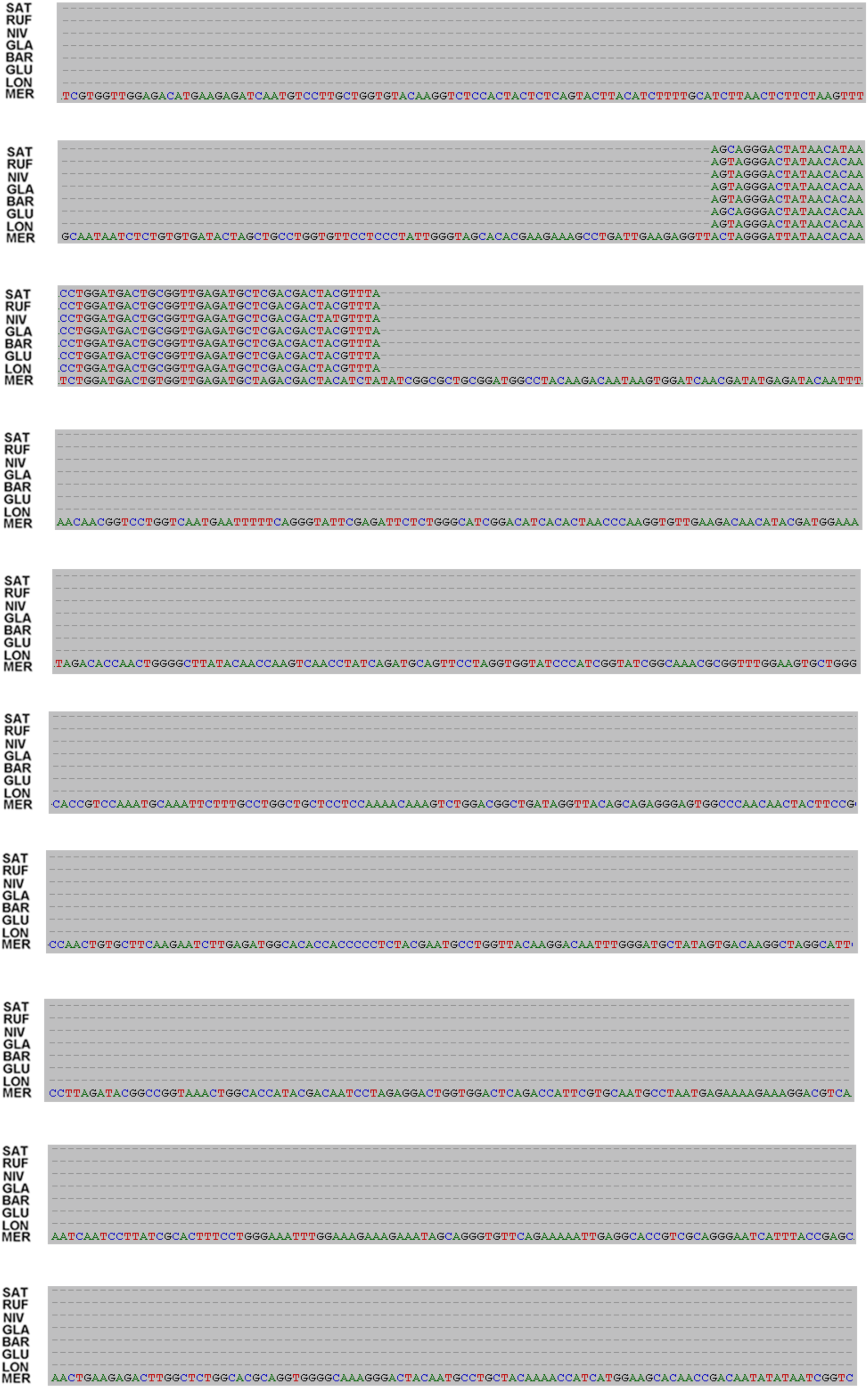
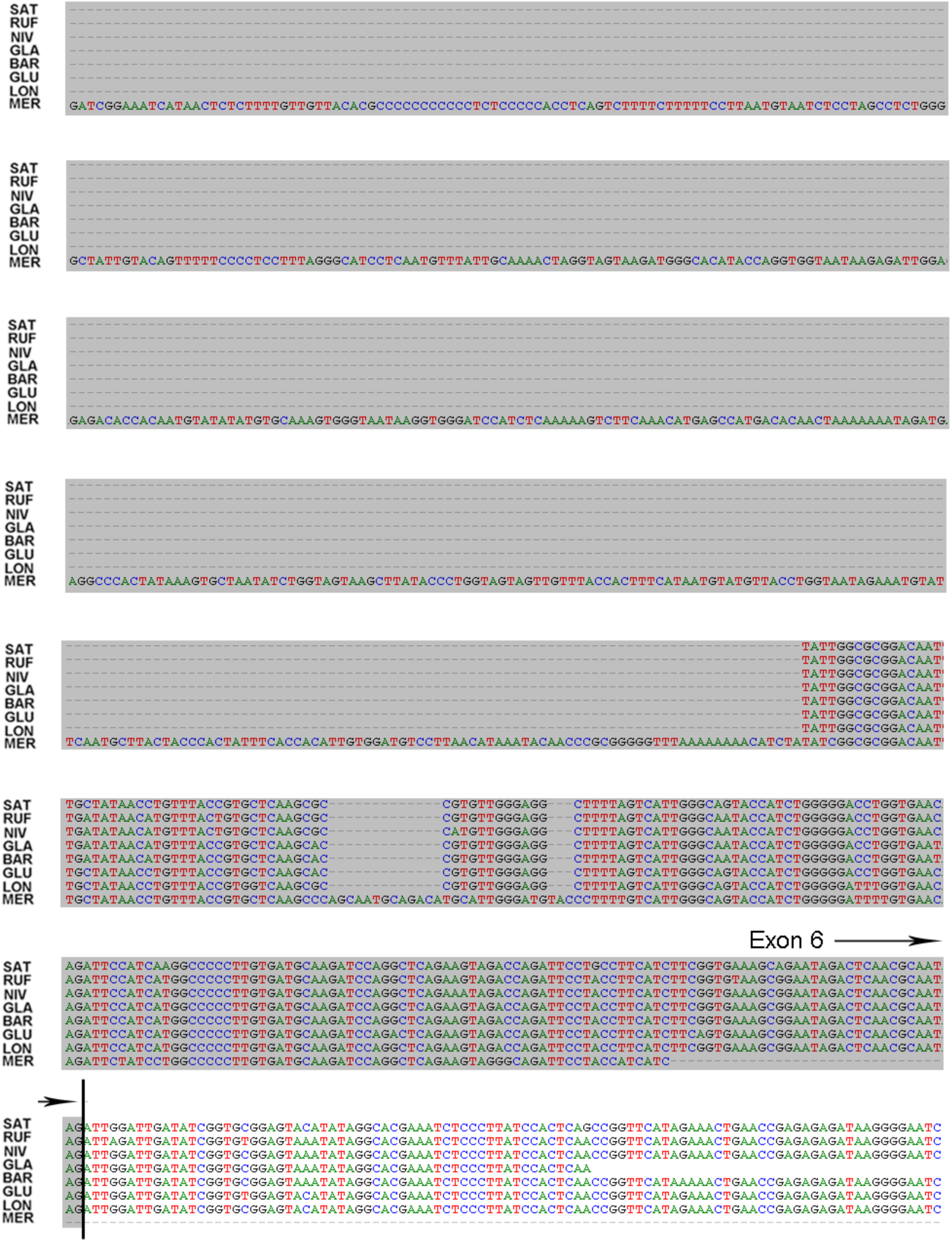
Sequence alignment of LOC_Os01g40320 orthologs across eight AA-genome *Oryza* species.

**Supplemental Figure 4.**
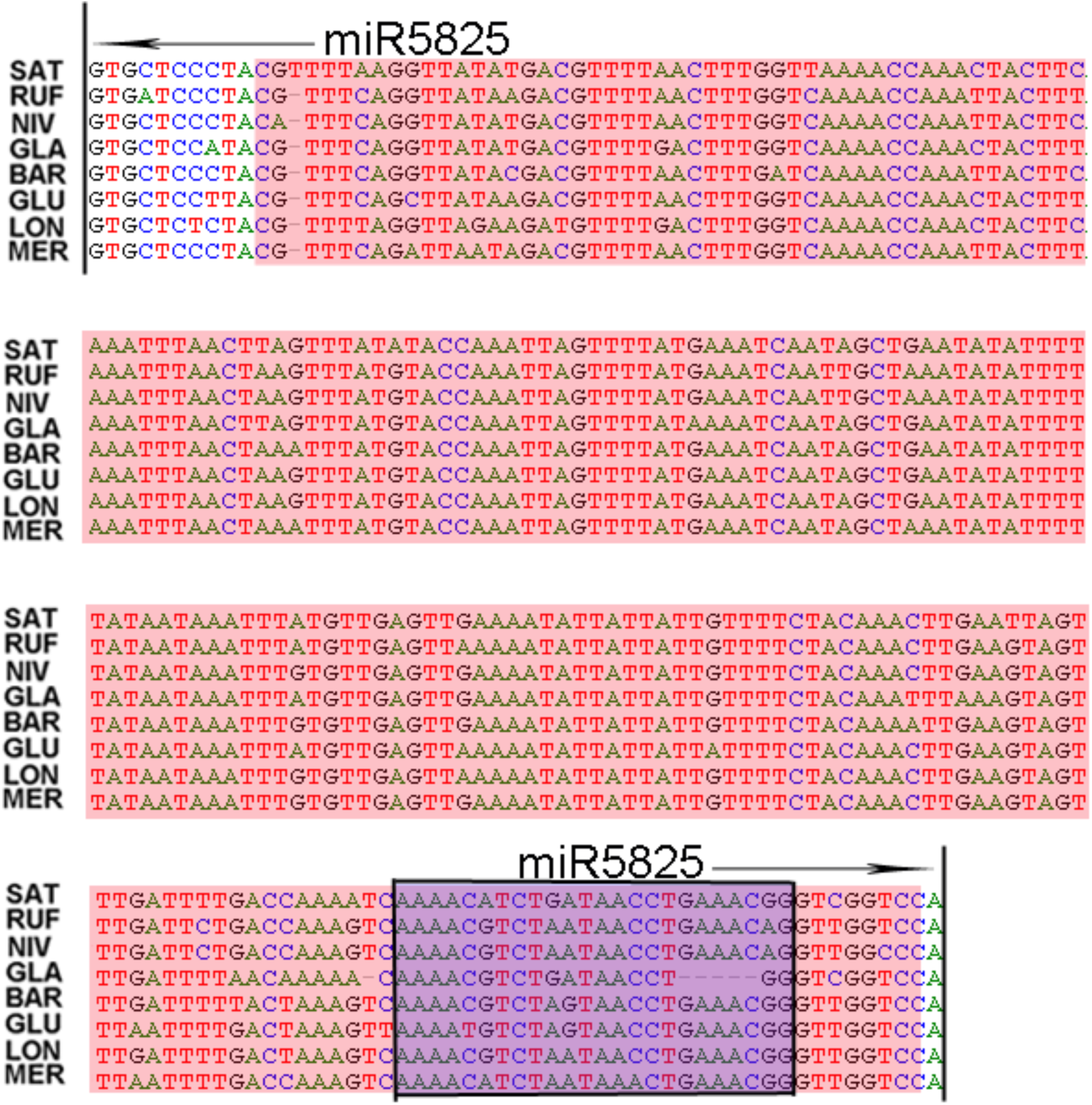
Alignment of osa-miR5825 orthologs across the eight AA-genome *Oryza* species. Sequences highlighted in red correspond to TEs (DNA transposon/*TcMar-Stowaway*); sequences in blue represent the mature miRNA within the MIRNAs.

**Supplemental Figure 5.**
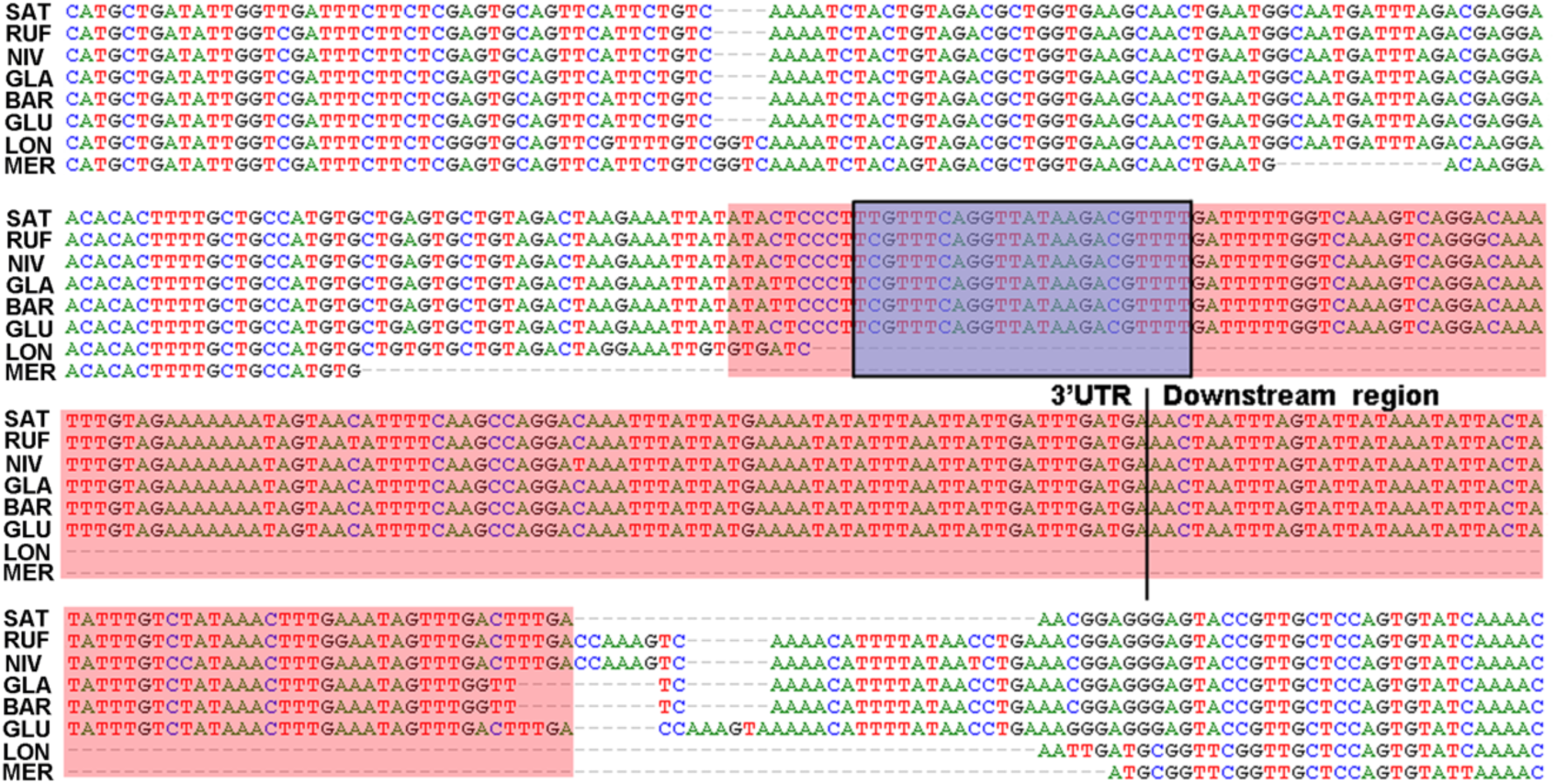
TE insertions leading to *de novo* generation of a miRNA target site in LOC_Os07g37680. The red box indicates the inserted TE sequence; the blue box marks the target site of osa-miR5825.

**Supplemental Figure 6.**
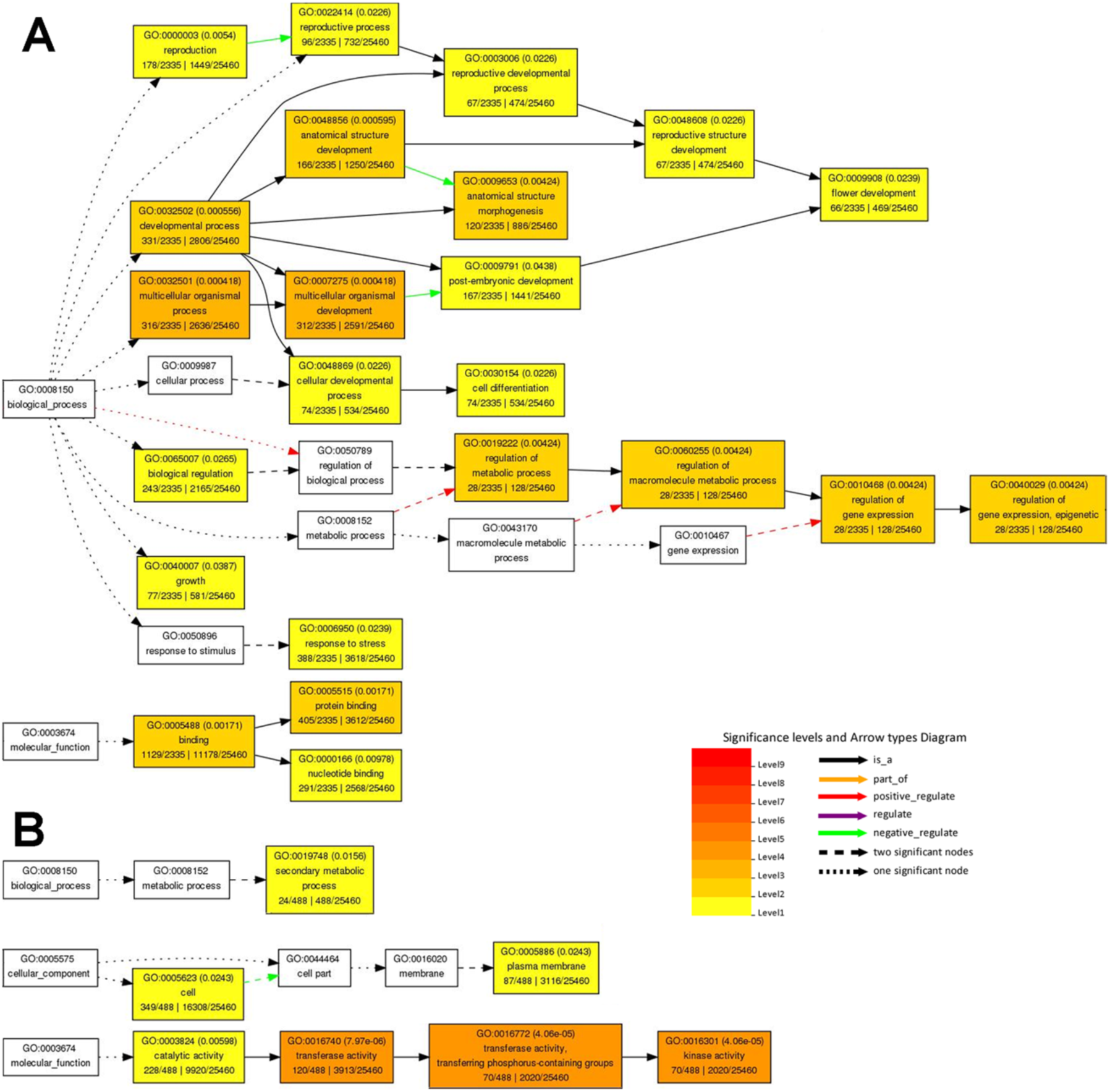
Gene Ontology (GO) enrichment analysis of miRNA target genes. (A) All miRNA target genes; (B) miRNA target genes whose target sites overlap TEs. The entire set of non-TE protein-coding genes from the SAT (*O. sativa*) genome was used as the background.

**Supplemental Figure 7.**
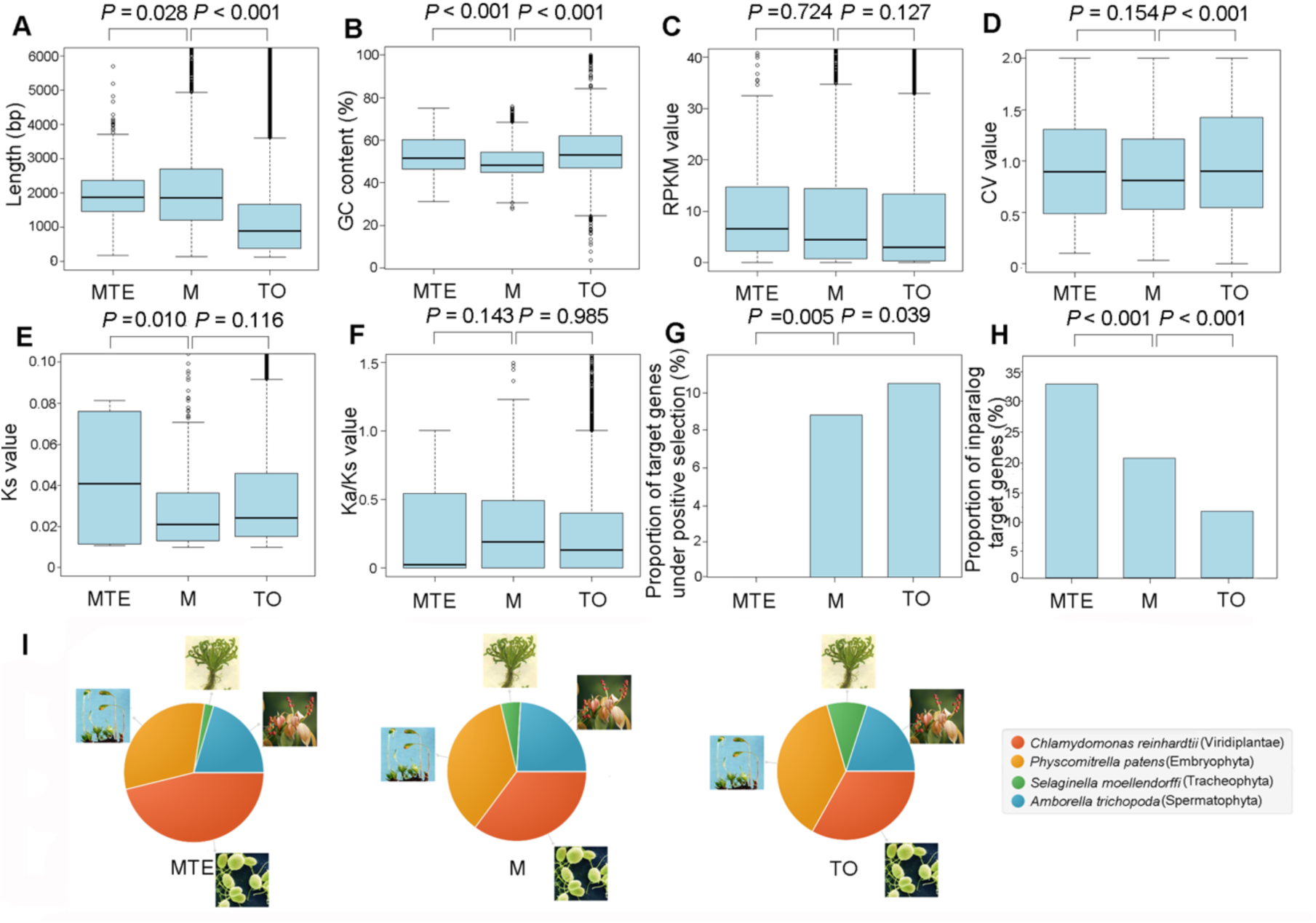
Comparison of features among miRNA target-gene categories. **(A)** Gene length; **(B)** GC content; **(C)** Gene expression level; **(D)** Gene expression variability; **(E)** *Ks*; **(F)** *Ka*/*Ks*; **(G)** Proportion of genes under positive selection; **(H)** Proportion of inparalog genes; **(I)** Distribution of evolutionary ages. MTE, miRNA target genes where both the MIRNA and its target site overlap the same TE; M, all miRNA target genes; TO, total genes. *P*-values in **(A)**–**(F)** were derived from permutation tests; *P*-values in (G) and (H) were obtained by chi-square tests.

**Supplemental Figure 8.**
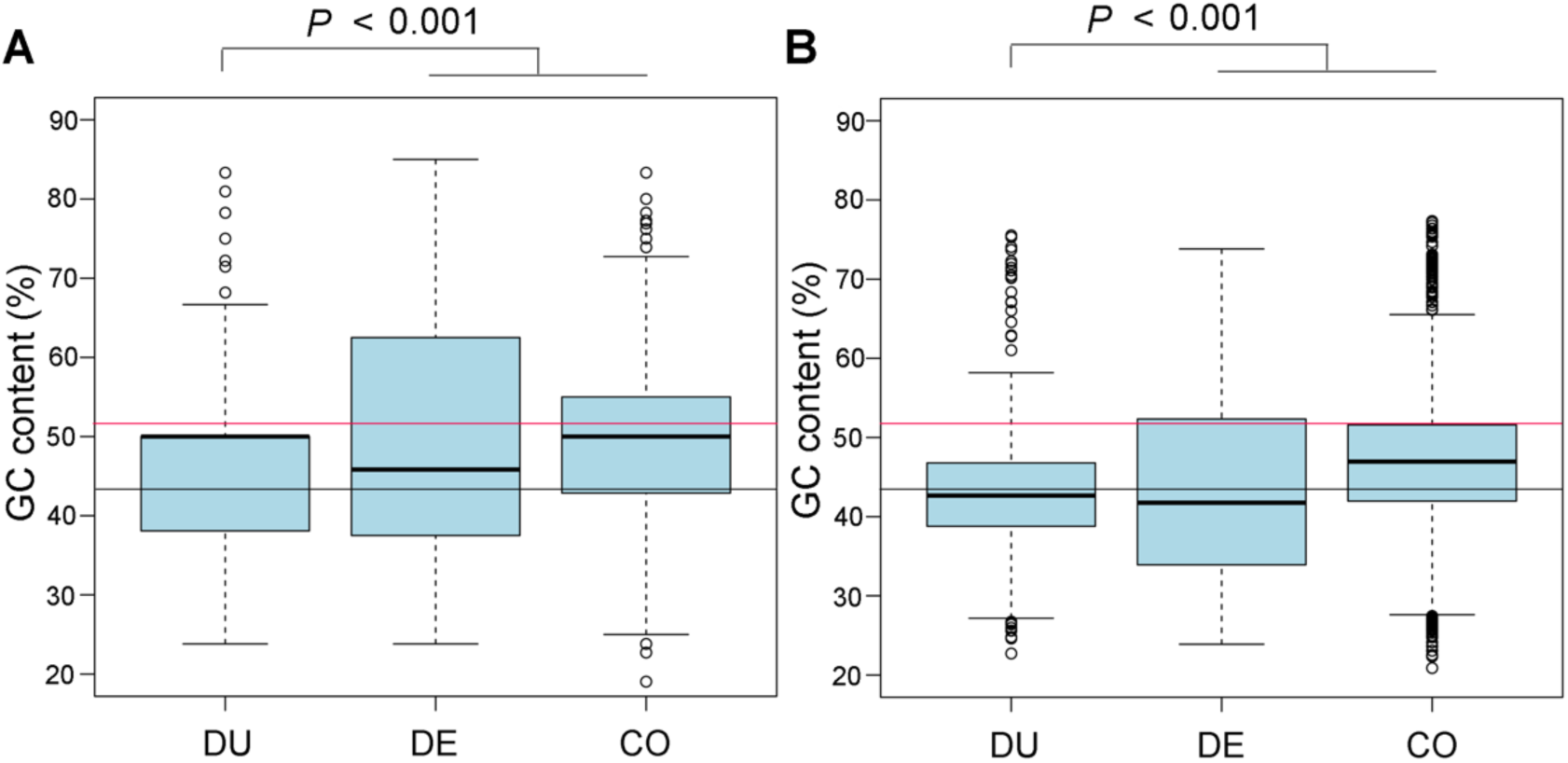
GC content of different MIRNA groups. **(A)** Mature miRNAs; **(B)** MIRNAs. DU, duplication-generated MIRNAs; DE, *de novo*generated MIRNAs; CO, conserved MIRNAs. The black horizontal line indicates the average genomic GC content of *O. sativa*; the red horizontal line shows the average GC content of *O. sativa* mRNAs. *P*-values were derived from permutation tests.

**Supplemental Figure 9.**
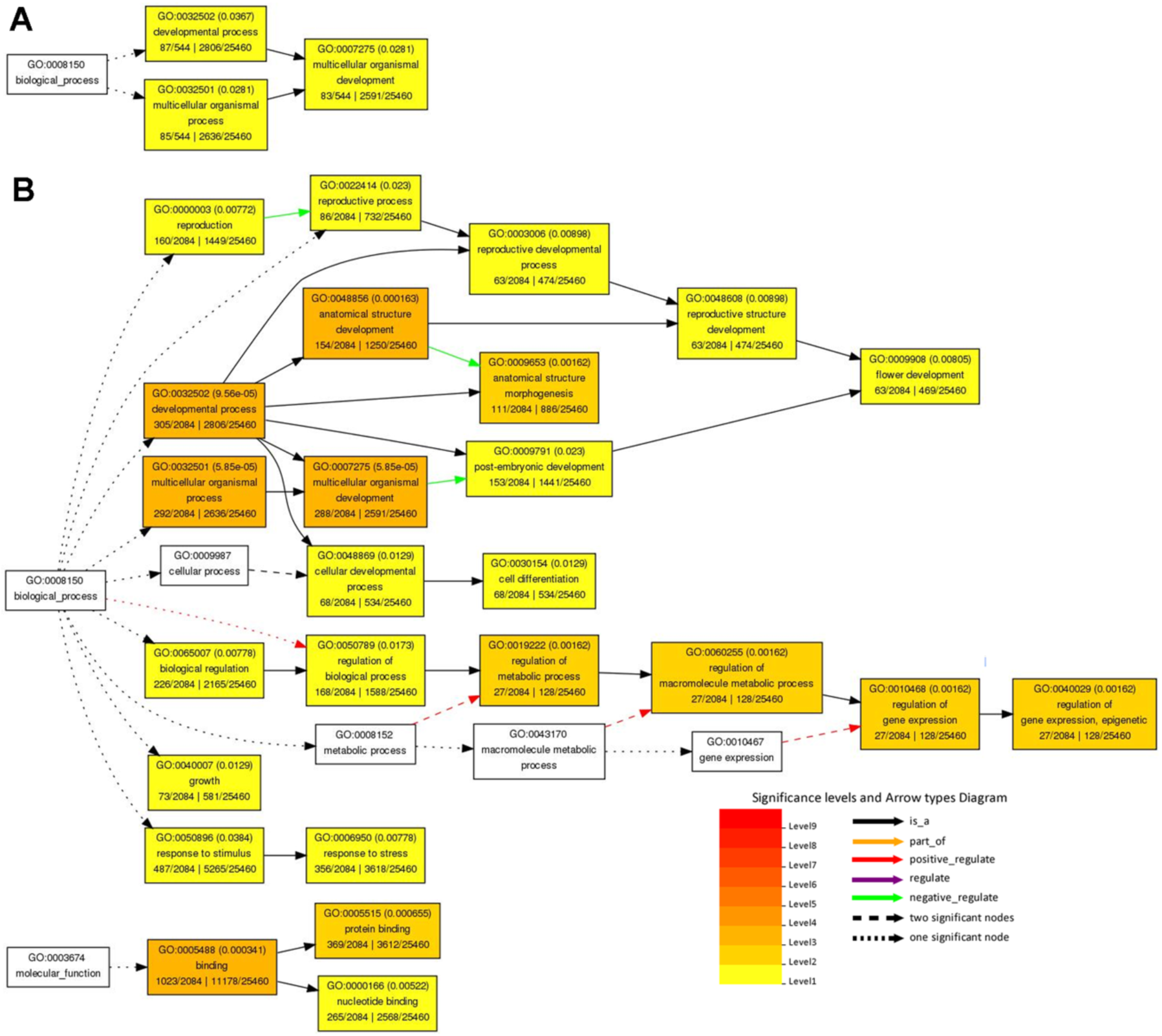
GO enrichment analysis of target genes from distinct MIRNA groups. **(A)** Target genes of duplication-generated MIRNAs; **(B)** Target genes of conserved MIRNAs. The entire set of non-TE protein-coding genes from the *O. sativa* genome served as the background.

### Supplemental Tables

**Supplemental Table 1. MIRNAs in the eight AA-genome *Oryza* species.** (see .xls file)

**Supplemental Table 2.** Counts of transposable elements overlapping MIRNAs in the eight AA-genome *Oryza* species.

|  | SAT | RUF | NIV | GLA | BAR | GLU | LON | MER |
| --- | --- | --- | --- | --- | --- | --- | --- | --- |
| **Class I :** |  |  |  |  |  |  |  |  |
| **LTR/Copia** | 1 | 1 | / | / | / | / | / | / |
| **LTR/Gypsy** | 2 | 5 | 2 | 4 | 5 | 2 | 3 | 2 |
| **Class II :** |  |  |  |  |  |  |  |  |
| **DNA** | 7 | 15 | 1 | 5 | 8 | 9 | 13 | 16 |
| **DNA/hAT** | 1 | 4 | 1 | 1 | 1 | 1 | 2 | 1 |
| **DNA/MuDR** | 4 | 5 | 7 | 6 | 7 | 3 | 3 | 4 |
| **DNA/TcMar-Stowaway** | 29 | 44 | 18 | 21 | 22 | 24 | 61 | 42 |
| **DNA/Tourist** | 6 | 6 | 4 | 5 | 5 | 5 | 7 | 7 |
| **RC/Helitron** | 6 | 5 | 3 | 3 | 4 | 2 | 3 | 4 |
| **Simple repeat:** | 6 | 8 | 4 | 4 | 4 | 7 | 3 | 4 |
| **Unknown:** | 14 | 14 | 8 | 12 | 12 | 13 | 6 | 11 |
| **Total** | 76 | 107 | 48 | 61 | 68 | 66 | 101 | 91 |

\* Number of MIRNAs overlapped with TEs in the eight AA- genome *Oryza* species were 62, 86, 41, 47, 53, 51, 76 and 69, respectively.
\*\* Number of MIRNAs in the eight AA- genome *Oryza* species were 310, 349, 273, 295, 296, 308, 340 and 326, respectively.

**Supplemental Table 3.** Counts of TEs overlapping miRNA target sites on mRNAs in the eight AA-genome *Oryza* species.

|  | SAT | RUF | NIV | GLA | BAR | GLU | LON | MER |
| --- | --- | --- | --- | --- | --- | --- | --- | --- |
| **Class I :** |  |  |  |  |  |  |  |  |
| **LTR** | 5 | 5 | 3 | 4 | 2 | 6 | 5 | 6 |
| **LTR/Copia** | 69 | 20 | 14 | 12 | 6 | 10 | 11 | 7 |
| **LTR/Gypsy** | 208 | 34 | 12 | 15 | 11 | 12 | 13 | 17 |
| **LINE** | 11 | 8 | 6 | 11 | 10 | 8 | 20 | 5 |
| **SINE** | 5 | / | / | 1 | / | / | 1 | 1 |
| **Class II :** |  |  |  |  |  |  |  |  |
| **DNA** | 12 | 20 | 7 | 2 | 4 | 11 | 1 | 7 |
| **DNA/hAT** | 1 | 3 | / | 1 | / | 2 | / | 5 |
| **DNA/MuDR** | 8 | 25 | 2 | 8 | 9 | 2 | 8 | 9 |
| **DNA/TcMar-Stowaway** | 86 | 453 | 89 | 70 | 178 | 52 | 474 | 188 |
| **DNA/Tourist** | 26 | 17 | 8 | 11 | 16 | 6 | 14 | 3 |
| **DNA/En-Spm** | 15 | 2 | / | 2 | 2 | / | 1 | / |
| **RC/Helitron** | 1 | 2 | 1 | 1 | / | / | 24 | 2 |
| **Simple repeat:** | 1 | 21 | 22 | 22 | 23 | / | 2 | 4 |
| **Unknown:** | 145 | 147 | 72 | 69 | 78 | 66 | 107 | 77 |
| **Total:** | 593 | 757 | 236 | 259 | 339 | 175 | 681 | 331 |

\* Numbers of MIRNAs which have target sites overlapped with TEs in the eight AA- genome *Oryza* species are 136, 132, 63, 69, 66, 56, 136 and 80, respectively.
\*\* Numbers of MIRNAs in the eight AA- genome *Oryza* species are 310, 349, 273, 295, 296, 308, 340 and 326, respectively.
\*\*\* Numbers of miRNA target genes on which miRNA target site overlapped with TEs in the eight AA- genome *Oryza* species are 311, 331, 159, 148, 188, 99, 231 and 155, respectively.
\*\*\*\* Numbers of miRNA target genes in the eight AA-genome *Oryza* species are 1016, 1419, 923, 864, 949, 738, 1141 and 914, respectively.

**Supplemental Table 4. Summary of MIRNAs whose sequences and target sites co-localize with the same TE.** (see .xls file)

**Supplemental Table 5.** Composition and classification of TEs in the genomic region surrounding LOC_Os01g40320 across the eight AA-genome *Oryza* species.

|  | SAT |  | RUF |  | NIV |  | GLA |  | BAR |  | GLU |  | LON |  | MER |  |
| --- | --- | --- | --- | --- | --- | --- | --- | --- | --- | --- | --- | --- | --- | --- | --- | --- |
|  | # | Length<br>(bp) | # | Length<br>(bp) | # | Length<br>(bp) | # | Length<br>(bp) | # | Length<br>(bp) | # | Length<br>(bp) | # | Length<br>(bp) | # | Length<br>(bp) |
| Class I : |  |  |  |  |  |  |  |  |  |  |  |  |  |  |  |  |
| LTR/Copia | 1 | 171 | 1 | 171 | 1 | 171 | 1 | 171 | 1 | 171 | 1 | 171 | 1 | 171 | / | / |
| LTR/Gypsy | / | / | / | / | / | / | 1 | 91 | / | / | / | / | / | / | / | / |
| LINE | 1 | 196 | 2 | 392 | 1 | 197 | 1 | 199 | 1 | 199 | 1 | 196 | 1 | 204 | 2 | 813 |
| SINE | / | / | / | / | 1 | 79 | 1 | 122 | 2 | 201 | 1 | 123 | 1 | 123 | / | / |
| Class II : |  |  |  |  |  |  |  |  |  |  |  |  |  |  |  |  |
|  | 3 | 523 | 2 | 342 | 3 | 523 | 3 | 524 | 3 | 524 | 3 | 515 | 5 | 642 | 5 | 811 |
| DNA | 6 | 1525 | 4 | 845 | 5 | 1151 | 6 | 1427 | 6 | 1253 | 5 | 1150 | 2 | 177 | 1 | 180 |
| DNA/MuDR | 2 | 1078 | 3 | 1197 | 4 | 1310 | 2 | 1087 | 3 | 941 | 2 | 1083 | 1 | 582 | / | / |
| DNA/hAT | 4 | 366 | 2 | 200 | 3 | 328 | 3 | 190 | 2 | 180 | 3 | 320 | 3 | 280 | 1 | 119 |
| DNA/TcMar | 3 | 561 | 5 | 843 | 2 | 462 | 2 | 488 | 2 | 488 | 2 | 507 | / | / | 2 | 458 |
| DNA/Tourist | 3 | 341 | 3 | 344 | 3 | 323 | 3 | 347 | 3 | 347 | 2 | 259 | 2 | 332 | 2 | 341 |
| Unknown: | 5 | 314 | 4 | 313 | 3 | 114 | 3 | 215 | 4 | 310 | 5 | 319 | 3 | 619 | 3 | 243 |
| Simple repeat: |  |  |  |  |  |  |  |  |  |  |  |  |  |  |  |  |

**Supplemental Table 6.** Sequence divergence of TEs overlapping osa-miR1868 and its target site in LOC_Os01g40320.

|  | **TEs overlapped with sat-miR1868 (%)** | **TEs overlapped with target sites of**<br>**osa-miR1868 on LOC_Os01g40320 (%)** |
| --- | --- | --- |
| SAT | 24.5 | 11.4* |
| RUF | 23.7 | 8.1 |
| NIV | 12.8 | 7.7 |
| GLA | 12.8 | 8.5 |
| BAR | 12.8 | 8.5 |
| GLU | 12.8 | 13.2* |
| LON | 12.8 | / |
| MER | 13.0 | / |

\* Compared with TEs overlapped with target sites of osa-miR1868 on LOC\_Os01g40320 in the RUF, NIV, GLA and BAR genomes, these two TEs showed elevated sequence divergence, suggesting that they underwent accelerated evolutionary divergences while evolving into exons;
\*\* Divergence value = mismatches of consensus sequence/(matches+mismatches).

**Supplemental Table 7.** Sequence divergence of TEs overlapping osa-miR5825 and its target site in LOC_Os07g37680.

|  | **TEs overlapped with osa-miR5825 (%)** | **TEs overlapped with target sites of**<br>**osa-miR5825 on LOC_Os07g37680 (%)** |
| --- | --- | --- |
| SAT | 13.8 | 5.0 |
| RUF | 13.1 | 5.5 |
| NIV | 12.9 | 6.3 |
| GLA | 16.2 | 5.9 |
| BAR | 14.3 | 5.9 |
| GLU | 14.7 | 5.5 |
| LON | 13.8 | / |
| MER | 14.3 | / |

\* Divergence value= mismatches of consensus sequence/(matches+mismatches)

**Supplemental Table 8.**
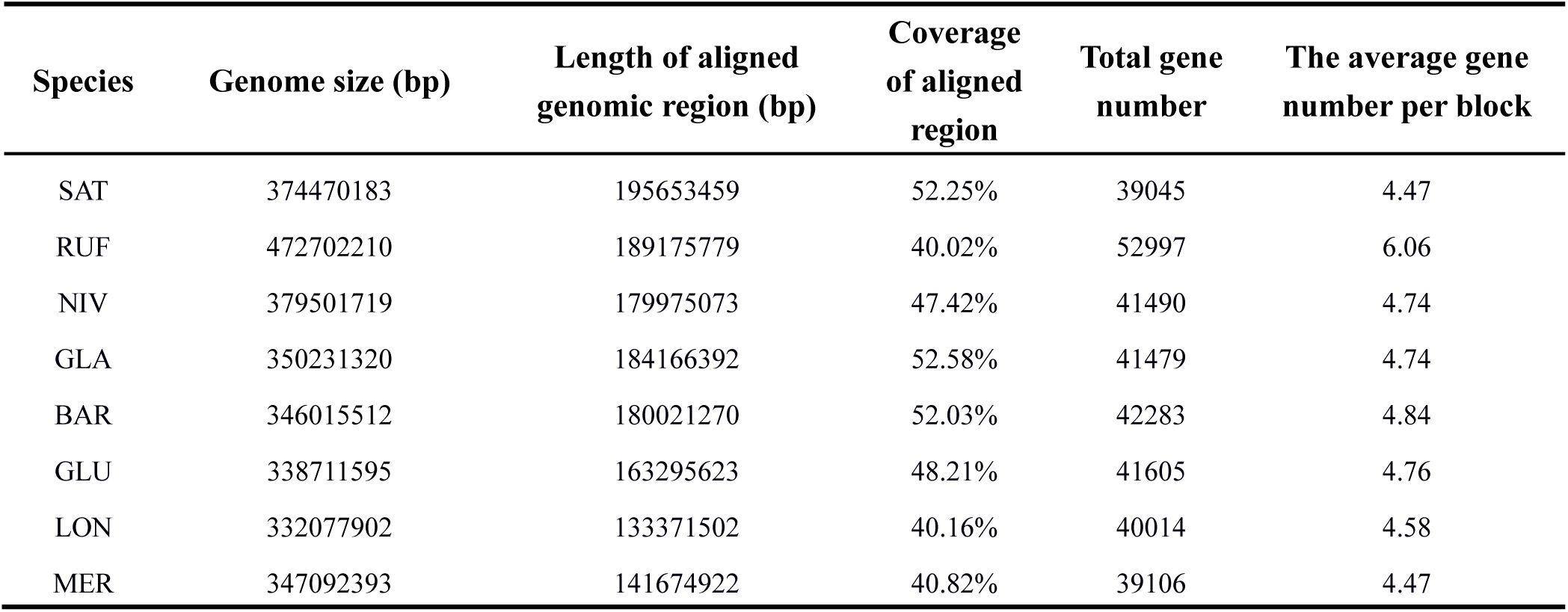
Summary of orthologous genomic regions across the eight AA-genome *Oryza* species.

